# Repurposing the evolutionary fitness of marine bacteria for water-conserving conversion of lignin-derived carbon

**DOI:** 10.1101/2025.05.20.655217

**Authors:** Ying Wei, Pei-Ru Chen, Bo Gong, Xian-Zheng Yuan, Shu-Guang Wang, Peng-Fei Xia

## Abstract

Reverting waste carbon into chemicals and materials with minimal freshwater reliance is essential for a sustainable future. Here, we report the establishment of a blue biological scheme by repurposing the evolutionary fitness of marine bacterium *Roseovarius nubinhibens* for efficient conversion of lignin-derived carbon into valuable chemicals with seawater as a water source. We created independency of the superior catalytic capacity of *R. nubinhibens* to circumvent a rare polar effect, and genetically decoupled the bioproduction pathway from central metabolism, channeling all lignin-derived carbon into desired products. Next, an orthogonal growth module was generated with acetate as a second carbon source, which is a ubiquitous byproduct and inhibitor in plant biomass hydrolysis, to sustain cell growth and supply necessary resources, but not substrates, for the growth-coupled bioproduction. Besides this designed metabolic segregation, we identified a cross-module push-and-pull synergy between bioproduction and cell growth, enabling enhanced performance and robustness. By deploying this biological scheme, we successfully achieved escalated conversion of the lignin-derived monomer 4-hydroxybenzoate to value-added compounds protocatechuate and β-ketoadipate with seawater, rather than freshwater, as the water source, generating a distinct blue biological scheme with potentials in simultaneous carbon and water conservation.

## Introduction

A sustainable future requires next-generation biotechnology to upcycle waste carbon and safeguard freshwater simultaneously. While carbon conservation has garnered increasing attention, the freshwater consumption of biosystems is often overlooked. Large amounts of freshwater is required as the solvent, medium or stabilizer in most biosystems, potentially competing with human demands, especially under the growing threat of regional and global water scarcity ^1,2^. One promising strategy is to leverage marine microbes and seawater to repurpose waste carbon into valuable chemicals with minimal freshwater reliance. Marine microbes often harbor unusual adaptive advantages and metabolic potentials, serving as ideal chassis to valorize diverse yet recalcitrant carbon compounds in waste streams, e.g., one-carbon compounds, alkanes and aromatics, to a broad-spectrum of chemicals ^3–7^. Seawater can guarantee a reliable water supply for biosystems, substituting a large amount of freshwater. Such blue biotechnology presents an innovative biological scheme for interdependent carbon and water conservation.

A universally longstanding challenge in reprogramming microbial chassis is to balance the conflict between engineered and natural behaviors, since microbes prioritize allocating available resources for survival and reproduction rather than bioproduction or other tasks designed for human needs ^8–10^. This is especially striking for microbes living in the ocean, where the oligotrophic environment forces them to assimilate and devote all accessible carbon to central metabolism. This evolutionary fitness becomes a biological barrier in engineering biology, preventing the directed reallocation of carbon flux towards bioproduction ^11^. Besides, the lagging development of synthetic biological toolsets makes metabolic reprogramming even more challenging compared to well-established microbial chassis. A previous report shows that engineered marine bacteria exhibit great promise in the degradation of lignin-derived monomers and can accumulate desired products by limiting the carbon flux towards central metabolism, while the products are metabolized eventually without harvesting in a certain production window ^12^.

Different approaches have been developed to tackle this intrinsic conflict. For instance, metabolic switches have been established based on biosensors, quorum sensing and optogenetic systems, allowing temporally metabolic shifts between growth and production ^13–17^. The core principle of these designs is to decouple bioproduction routes from the central metabolic network, channeling maximal carbon towards designed products in a defined period. However, bioproduction demands on enzymes, cofactors, and commonly energy and reducing power, thereby asking for efficient turnovers of functional proteins and redox intermediates, e.g., ATP and NAD(P)H, which can hardly be independent from cell growth ^18,19^. One alternative is to genetically separate the bioproduction pathway and central metabolism, generating two orthogonal yet synergistic modules ^20^. These two modules can be fueled by two carbon sources: one solely serves as the substrate for bioproduction, while the other independently supports cell growth, providing necessary resources but not the substrate for bioproduction.

Lignin constitutes 15 - 40% of the plant biomass on Earth, giving an important but recalcitrant carbon source for valorization ^21–23^. Previous studies engineered *Pseudomonas putida*, a soil bacterium with diverse metabolic potentials, to convert lignin-derived carbon to value-added chemicals (e.g., β-ketoadipic acid and muconic acid) with glucose or part of the lignin monomer as a second carbon source for cell growth, demonstrating the potential for modular reprogramming ^24–26^. However, glucose is an immediately fermentable sugar in the hydrolysate of plant biomass, which can be directly converted by most model chassis (e.g., *Saccharomyces cerevisiae*) and may lead to food-related debates at large scale. A preferred but unexplored alternative is the byproducts from the pretreatment of plant biomass, such as acetate, to power cell growth and provide necessary resources for bioproduction from lignin-derived carbon. This new scheme generates a win-win scenario for upcycling waste carbon and balancing engineered and natural microbial metabolism.

Here, we report the water-conserving conversion of lignin-derived carbon into value-added compounds through modularly reprogramming marine *Roseobacter clade* bacterium *Roseovarius nubinhibens*. First, we determined the catalytic capability of 4-hydroxybenzoate 3-monooxygenase (PobA) from *R. nubinhibens* and generated a bioproduction module by creating the independency of *pobA* and decoupling the bioproduction pathway from central metabolism. Next, a growth module was formed by co-feeding acetate, a ubiquitous byproduct and inhibitor in plant biomass hydrolysis, sustaining cell growth and supplying resources for bioproduction. In addition, the isotopic labeling and metabolomics analysis showed synergistic interactions between these genetically independent modules. Leveraging the reprogrammed *R. nubinhibens*, the lignin-derived monomer 4-hydroxybenzoate (4HB) was efficiently valorized to pharmaceutical compound protocatechuate (PCA) with seawater as the water source. The strategy was further extended from a single-step enzymatic conversion to pathway-level modulation for producing β-ketoadipate (β-KA), a promising precursor for performance-advantaged bioproducts, underscoring its potential in simultaneous carbon and water conservation.

## Results

### Catalytic advantage of the PobA from *R. nubinhibens* under high salinity and pH

The flavin-dependent monooxygenase PobA, one of the 4-hydroxybenzoate-3-hydroxylases (PHBHs), plays an indispensable role in the biological valorization of lignin-derived carbon, particularly the fundamental 4-hydroxyphenyl units of depolymerized lignin ^27–29^. Specifically, PobA catalyzes the hydroxylation of 4HB, a lignin-derived aromatic compound, to PCA, a pharmaceutical monomer and the key metabolic node connecting to various platform compounds, such as acetyl-CoA through the native β-ketoadipate pathway, pyruvate through gluconeogenesis, and catechol from synthetic pathways, for the production of high value-added chemicals, such as lactate and muconic acids **(Fig. 1A)** ^30–32^. However, the hydroxylation is considered as a biological bottleneck in the β-ketoadipate pathway. Given the advances in synthetic biology, elevating the activity of PobA has been the primary strategy to engineer microbes for efficiently upcycling lignin-derived monomers ^33–35^.

**Fig. 1.**
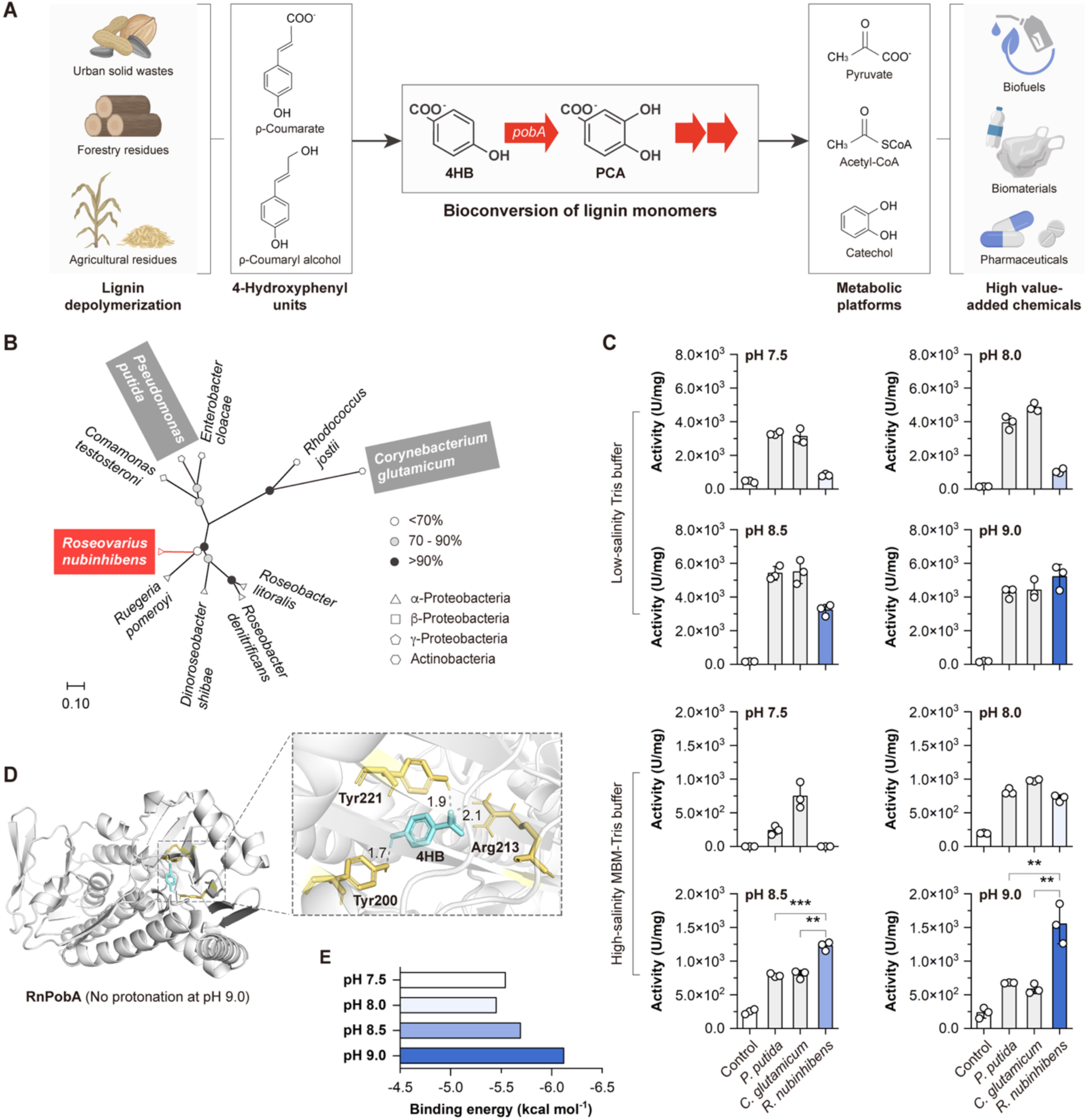
The catalytic potential of PobA from *R. nubinhibens*. **(A)** Schematic illustration of bioconversion of lignin monomers with *pobA* as an essential step. Through this process, 4-hydroxyphenyl units of depolymerized lignin can be converted to high valued-added chemicals via various platform compounds. **(B)** Phylogenetic analysis of PobA from *R. nubinhibens* and other organisms. The phylogenetic tree was built using the maximum likelihood method. Bootstrap values generated by 1,000 replicates are indicated at branch nodes with black circles representing >90%, grey circles representing 70 - 90% and white circle representing <70%. The taxonomy is indicated at branch terminals with the triangle representing α-Proteobacteria, the square representing β-Proteobacteria, the pentagon representing γ Proteobacteria and the hexagon representing Actinobacteria. The scale bar indicates evolutionary distances. **(C)** Enzyme activity of PobA. The *pobA* genes were expressed in *R. nubinhibens* and the enzymes were extracted and analyzed in low salinity Tris and high-salinity MBM-Tris buffer. The experiments were conducted in triplicate, and the circles and error bars represent the individual values and standard deviations of three biological replicates, respectively. The differences were statistically evaluated by *t*-test (**, P<0.01; ***, P<0.001, unpaired and two-tailed). **(D)** Molecular docking result of RnPobA with 4HB at pH 9.0. The residues of RnPobA identified as the binding site to 4HB are indicated in yellow and the hydrogen bonds are indicated in grey. **(E)** Binding energy of RnPobA with 4HB at different pHs. Panel **A** created with BioRender.com.

Marine *R. nubinhibens,* an abundant bacterium belongs to the *Roseobacter* clade in coastal areas, has shown unusual accumulation of PCA when growing on 4HB ^12^. This mainly resulted from a potential imbalance between PobA and downstream PcaHG, which has been reported as a bottleneck in 4HB metabolism ^36^. We have taken advantage of this feature by amplifying the imbalance between PobA and PcaHG for enhanced PCA production ^12^. Beyond production, the accumulation of PCA suggests a salt-tolerant PobA in *R. nubinhibens,* which may be harnessed for water-conservative bioconversions. In a preliminary comparison, *R. nubinhibens* shows a similar substrate utilization rate but better product accumulation of PCA compared to *P. putida*, the well-established chassis for efficient lignin valorization, under high salinity and pH **(Fig. S1)**. However, the catalytic capacity of this marine PobA remains to be explored, especially when compared to that of the rising lignin-valorizing chassis *P. putida* and *Corynebacterium glutamicum*. Through phylogenetic analysis of PobA, the marine *Roseobacter* clade bacterium *R. nubinhibens,* as α-Proteobacteria, is evolutionarily distinct from *P. putida* and *C. glutamicum*, which belong to γ-Proteobacteria and Actinobacteria, respectively **(Fig. 1B)**. This distant evolutionary relationship suggests a unique fitness of *R. nubinhibens* adapting to the ocean. It should be noted that the *pobA* gene is located in the *pca* operon, and all these genes are regulated by the PcaQ transcription activator located upstream of this operon in *R. nubinhibens* ^37^. While key genes in the operon, such as *pobA*, *pcaH*, and *pcaG*, in *R. nubinhibens* and *P. putida* share 60 - 80% of the percent identities **(Table S1)**, the genomic arrangements are quite different. In *P. putida*, the *pobA* gene (PP_RS18385) is regulated by PobR and separated from *pcaHG* (PP_RS24255 and PP_RS24250) ^38^.

To quantitatively characterize PobA, three *pobA* genes from *P. putida* (PpPobA), *C. glutamicum* (CgPobA) and *R. nubinhibens* (RnPobA) were expressed on plasmids in the model strain *Escherichia coli* BW25113 and *R. nubinhibens* at different salinities and pHs **(Fig. S2 and Table S2)**. We used low-salinity Tris buffer and high-salinity MBM-Tris buffer with higher concentration of NaCl and MgSO_4_ to test the enzyme activity **(Table S3)**. Though the main salinity of both buffers is lower than seawater, we found that the PobA activities, despite of the origins, were significantly impacted by salinities, while RnPobA was less sensitive to higher salinity in the two hosts. At pH 8.0 and 8.5, PpPobA and CgPobA reached comparably maximal activities, while the activities were repressed at pH 9.0 **(Figs. S3 and S4)**. Differently, the activity of marine RnPobA raised along with the increasing pH, although it was lower than that of PpPobA and CgPobA under low salinity at pH 7.5-8.5 and under high salinity at pH 7.5-8.0. Interestingly, RnPobA found its unique niche under high salinity and high pH, reaching the maximal catalytic activity of 1224.85 ± 52.78 U/mg and 1552.33 ± 240.91 U/mg with the high-salinity MBM-Tris buffer at pH 8.5 and 9.0, respectively, which was significantly higher than PpPobA (789.06 ± 20.93 U/mg and 677.02 ± 5.48 U/mg) and CgPobA (801.57± 49.63 U/mg and 585.60 ± 56.95 U/mg) **(Fig. 1C)**. RnPobA also showed host-dependent activity with a preference of *R. nubinhibens* over *E. coli* **(Figs. S3 and S4)**, indicating a distinct hydroxylation capacity of RnPobA in marine chassis under conditions mimicking the ocean. While some PHBHs, such as PpPobA, exhibit specificity to NADPH, PraI from *Paenibacillus* sp. JJ-1b shows cofactor preference to both NADH and NADPH ^35^. Due to the importance of cofactors in bioconversion, we determined the cofactor preference of RnPobA and discovered that the relative enzyme activity of RnPobA reached 2272.59 ± 182.79 U/mg with NADPH and only 266.18 ± 110.70 U/mg with NADH at pH 8.5 **(Fig. S5)**, showcasing its preference to NADPH over NADH.

Moreover, we deciphered the underlying mechanism of the catalytic property of RnPobA under high salinity in silico. The protein structures were pretreated at varied pHs predicted by AlphaFold. The results illustrated that RnPobA changed its proton numbers in response to different pHs, while PpPobA lost the protons at pH 8.0 to 9.0 and CgPobA held the same proton number **(Table S4 and Fig. S6)**. Through molecular docking, we found that the binding sites of RnPobA for 4HB hydroxylation were Tyr200, Tyr221 and Arg213 residues at pH 9.0, and Pro292 and Arg213 residues at pH 7.5, 8.0 and 8.5, differing from that of PpPobA (Thr103, Arg44, Arg214 and Arg218) and CgPobA (Tyr202, Tyr223 and Arg215) **(Figs. 1D and S7)**. These changes eventually led to varied hydrogen bonds and decreased binding energies between RnPobA and 4HB from -5.54 to -6.12 kcal/mol when pH increased from 7.5 to 9.0 **(Fig. 1E)**. To the contrary, the binding energies of PpPobA and CgPobA were maintained around -5.86 and -5.85 kcal/mol **(Fig. S8)**.

### Rescuing 4HB hydroxylation with independent RnPobA

We attempted to circumvent the biological bottleneck of 4HB hydroxylation under conditions simulating seawater by leveraging RnPobA in its native host *R. nubinhibens* **(Fig. 2A)**. By introducing early STOP codons with CRISPR-Cas base editing, we deactivated the *pcaH* and *pcaG* genes, located the downstream of *pobA* and encoding the protocatechuate 3,4-dioxygenase β-subunit (PcaH) and α-subunit (PcaG), to block the PCA cleavage pathway and decouple bioproduction from central metabolism for product accumulation **(Fig. 2A)**. The resulting *pcaH*-deficient WY06 (*pcaH* Gln40*) and *pcaG*-deficient WY07 (*pcaG* Gln3*) **(Table S5)** lost the ability to convert 4HB to PCA ^37^. This was reasonable that, in these two strains, 4HB, as the sole carbon source, could not enter the central metabolism to sustain cell growth and provide necessary resources for PCA production. We assumed that the accumulation of enzymes, cofactors and energy in advance could recover the conversion of 4HB. To validate the assumption, the wild-type and WY06 strains were cultivated in Marine Broth 2216 (MB2216) liquid media until late exponential phase, and then, the cells were concentrated in fresh marine basal media (MBM) to an initial OD_600_ of 5.0, followed by feeding 4HB as the substrate. Surprisingly, the *pcaH*-deficient WY06 still failed to convert 4HB to PCA, while the concentrated wild-type strain remained the functionality with a PCA yield at 16.97 ± 3.24% **(Fig. S9 and Table S6)**. These results implied that the *pobA* gene was influenced by the deactivated genes located its downstream of the same operon due to a potentially rare polar effect. We complemented the *pcaH* and *pcaG* genes in the deficient strains WY06 and WY07, respectively, but still substantially interfered growth was observed, demonstrating the unexpected polar effect **(Fig. 2B)**. While polar effects usually impair downstream genes caused by the disruption of an upstream locus ^39^, we notice two similar observations of such rare events in two *E. coli* operons ^40,41^. Additionally, we identified dramatically lower mRNA levels of *pobA* with 98.82%- and 98.73%-decrease in *pcaH*- and *pcaG*-deficient strains, respectively, compared to the wild-type strain, exhibiting strong transcriptional disruptions of this polar effect **(Fig. 2B)**.

**Fig. 2.**
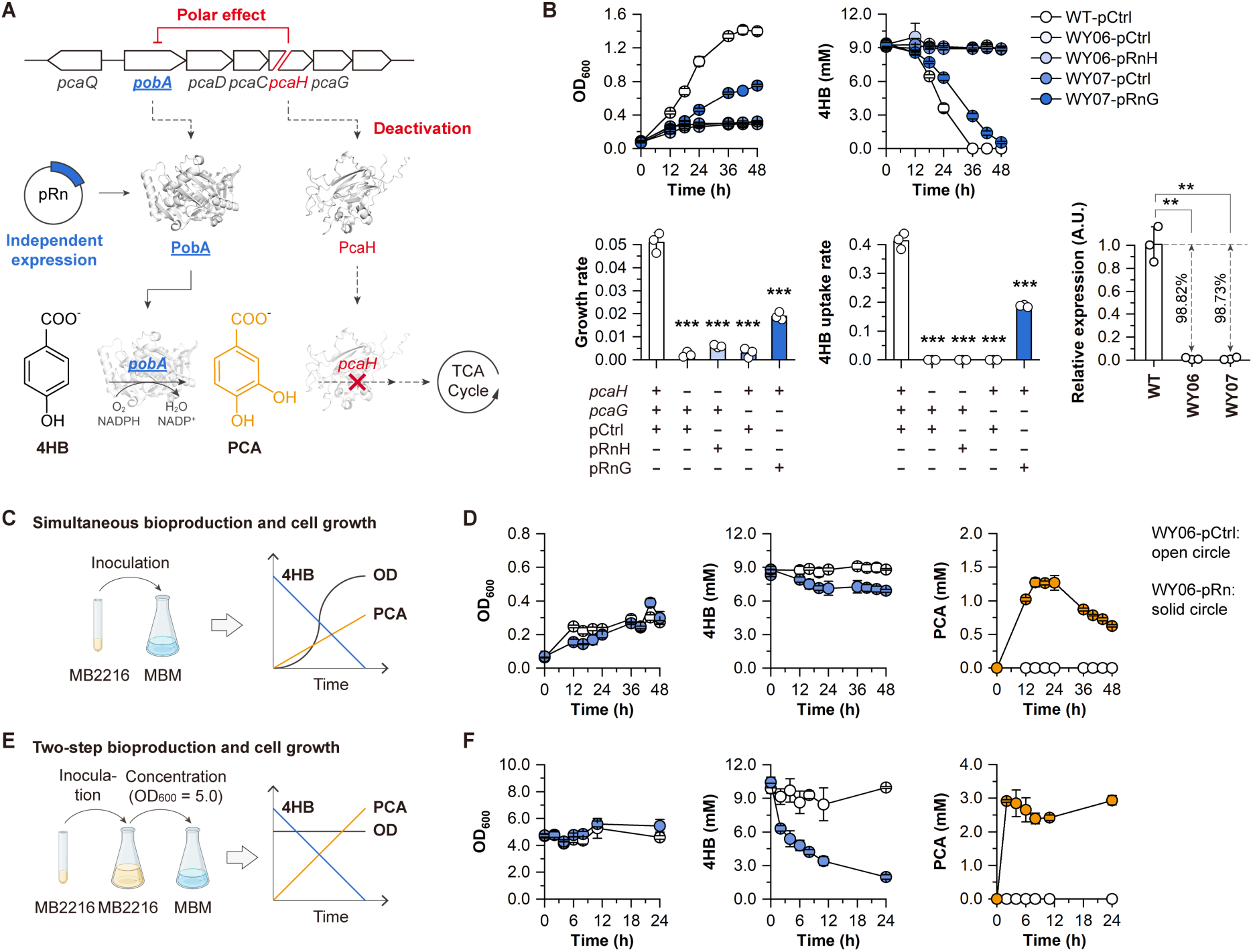
Rescue of the hydroxylation from 4HB to PCA. **(A)** Designed PCA bioproduction module from 4HB. The *pobA* gene was independently expressed and *pcaH* was deactivated. **(B)** Validation of the rare polar effect via growth and substrate uptake and transcription levels of *pobA* in WY06 and WY07. The wild-type strain with pCtrl (WT-pCtrl) and the strains WY06-pCtrl and WY07-pCtrl served as controls. The differences were statistically evaluated by *t*-test (**, P<0.01; ***, P<0.001, unpaired and two-tailed). **(C)** Design principle and **(D)** bioproduction performance of the WY06-pRn with simultaneous bioproduction and cell growth. The WY06-pCtrl and WY06-pRn strains were first cultivated in MB2216 overnight and then inoculated into MBM with 4HB as the sole carbon source with a dilution rate of 1:50. **(E)** Design principle and **(F)** bioproduction performance of the WY06-pRn with a two-step of bioproduction and cell growth. The WY06-pCtrl and WY06-pRn strains were first cultivated in MB2216 overnight and then inoculated into MB2216 with a dilution rate of 1:50. After cultivation overnight, cells were concentrated in MBM to an initial OD_600_ of 5.0, followed by feeding 4HB as the sole carbon source. Samples were taken at different time intervals. The experiments were conducted in triplicate, and the circles and error bars represent the individual values and standard deviations of three biological replicates, respectively. Panels **C** and **D** created with BioRender.com.

To enable the conversion from 4HB to PCA, the *pobA* gene needs to be released from the restriction of polar effect. According to the explored mechanism, one strategy is to introduce catalytic-dead point mutations or in-frame deletions, which inactivates a gene at post-translational level, to avoid the impacts of polar effect on mRNAs. Yet, this strategy requires in-depth mechanical information of RnPobA, which is currently unavailable. Therefore, we designed to create gene independency of *pobA* by delivering the *pobA*-expressing plasmid pRn into WY06 **(Tables S2 and S5)**. The resulting WY06-pRn successfully consumed 1.65 ± 0.17 mM 4HB and produced 1.26 ± 0.02 mM PCA **(Fig. 2C, 2D and Table S6)**. The molar yields of PCA reached 77% regardless of initial substrate levels, which were significantly higher than concentrated wild-type strain **(Figs. S9 and S10)**. These results illustrated that the independent expression of *pobA* rescued the hydroxylation process from 4HB to PCA in the operon-disrupted strain. However, PCA production could only be observed with cell growth on yeast extract in the minimal media **(Fig. 2D)**, which was indicated by the similarly marginal growth with different initial concentrations of 4HB. When the cells stopped growing at the final OD_600_ of 0.3, PCA production was terminated as well, implying its dependence on cell growth. To relieve this constraint, we concentrated the grown cells to an OD_600_ of 5.0 before supplying 4HB for PCA production **(Fig. 2E)**. WY06-pRn converted 4.10 ± 0.49 mM 4HB to 2.91 ± 0.03 mM PCA in 2 h, exhibiting a 2.31-fold rise in titer while retaining comparably high yield (71.96 ± 9.40%) **(Fig. 2F and Table S6)**. Thereby, we successfully established a bioproduction module leveraging RnPobA and reprogrammed marine bacteria to channel maximal 4HB to PCA, while the growth-dependent resources have to be complemented in grown cells through a stepwise strategy that maintains the decoupled bioproduction and cell growth.

### Carbon co-utilization drives the orthogonal bioproduction and cell growth

We introduced a second carbon source to sustain cell growth and continuously provide necessary resources and reducing power for bioproduction. Acetate is a universal byproduct from plant biomass hydrolysis and an inhibitor of conventional fermentation processes ^42,43^. By utilizing acetate as the additional carbon source, two forms of waste carbon from plant biomass can be upcycled concurrently, relieving a common stress on fermentation and avoiding competition for immediately fermentable carbon (e.g., glucose). Notably, *R. nubinhibens* cannot utilize common sugars, such as glucose and xylose **(Fig. S11)**, and therefore does not compete for these carbon sources even when they are present. We found that *R. nubinhibens* could grow on acetate and the OD_600_ reached 0.75 **(Fig. S12)**, showcasing the possibility of introducing acetate as the second carbon source to form the growth module. By co-feeding different carbons to the genetically decoupled bioproduction and growth modules, we conceived that 4HB can be channeled solely for PCA production, while acetate supports cell growth and supplies necessary resources for bioproduction **(Fig. 3A)**.

**Fig. 3.**
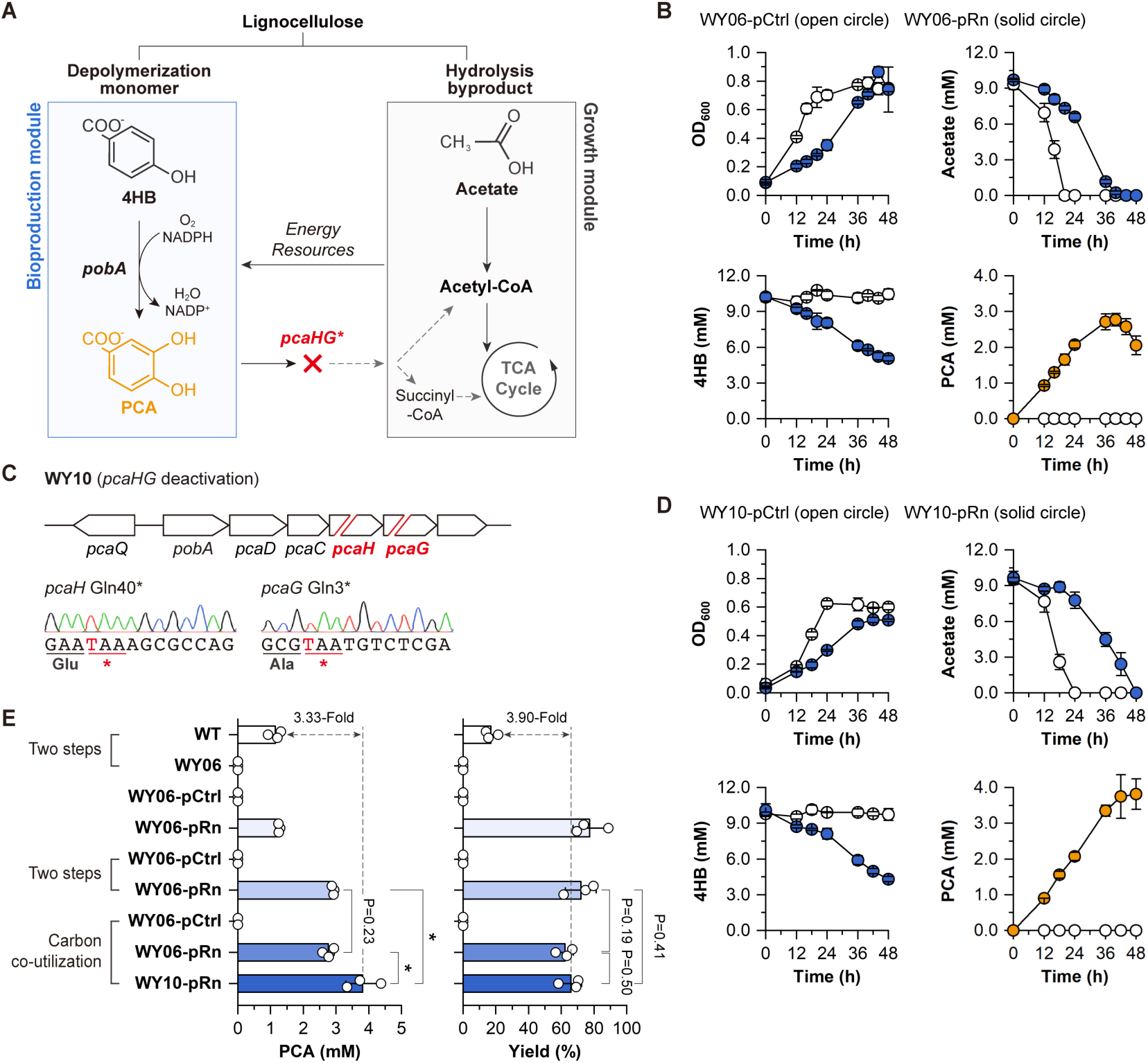
Modularly reprogrammed bioproduction and cell growth. **(A)** Design scheme of the bioproduction and cell growth modules. In the bioproduction module, 4HB was converted to PCA, and in the growth module, acetate was utilized to sustain cell growth and supply resources for bioproduction. **(B)** Bioproduction performance of the WY06-pRn with two carbon sources. The WY06-pCtrl and WY06-pRn strains were cultivated in MBM with 4HB and acetate. **(C)** Genome editing results of *pcaHG* in WY10. Based on WY06, the *pcaG* gene was deactivated with base editing by introducing an early STOP codon, generating WY10 (*pcaH* Gln40* and *pcaG* Gln3*). The introduced STOP codons are indicated in red. **(D)** Bioproduction performance of the WY10-pRn with two carbon sources. The WY10-pCtrl and WY10-pRn strains were cultivated in MBM with 4HB and acetate. **(E)** Titers and molar yields of PCA from 4HB. Samples were taken at different time intervals. The experiments were conducted in triplicate, and the circles and error bars represent the individual values and standard deviations of three biological replicates, respectively. The differences were statistically evaluated by *t*-test (*, P<0.05, unpaired and two-tailed).

Acetate and 4HB were simultaneously utilized to cultivate the reprogrammed *R. nubinhibens* strain WY06-pRn. While WY06-pRn displayed slower growth and acetate consumption rates, its final OD_600_ was similar to WY06-pCtrl (WY06 with a control plasmid) **(Fig. 3B)**. WY06-pRn converted 4.46 ± 0.21 mM 4HB to 2.77 ± 0.14 mM PCA with the yield of 62.25 ± 5.21% **(Fig. 3B and Table S6)**, which was equivalent to the stepwise strategy using concentrated WY06-pRn **(Fig. 2D)**, demonstrating that the growth module driven by acetate could enable the bioproduction of PCA from 4HB. However, we found that the produced PCA was metabolized after 40 h, which was similar to previous reports on engineered *P. putida* ^25,44^. According to previous work in other strains, we hypothesized that the remaining α-subunit of protocatechuate 3,4-dioxygenase (encoded by *pcaG*) in WY06 reserved certain ring-cleavage activity for PCA. To resolve this issue, we employed the single-nucleotide resolution CRISPR-Cas base editing system to deactivate *pcaG* by introducing an early STOP codon to *pcaG* in WY06 and generated WY10 with deactivated *pcaH* (Gln40*) and *pcaG* (Gln3*) **(Fig. 3C and Table S5)**. We delivered the *pobA*-expressing plasmid pRn and the control plasmid pCtrl into WY10, giving WY10-pRn and WY10-pCtrl, respectively. As a result, WY10-pRn produced 3.81 ± 0.43 mM PCA with a molar yield at 65.85 ± 5.47%, exhibiting another 1.38-fold increase in titer over WY06-pRn without further degradation **(Fig. 3D and Table S6)**. Despite the endeavors, the molar yields of PCA remained below the theoretical maximum. We hypothesized three routes of the loss of carbon. First, 4HB might react with all-trans-polyprenyl diphosphate, catalyzed by the 4-hydroxybenzoate octaprenyltransferase (EC 2.5.1.39) encoded by the *ubiA* gene in *R. nubinhibens* (ISM_RS05895) ^45^, thereby reducing the PCA yield. Second, the intracellular and extracellular distribution of PCA might reduce its yield. Yet, we did not detect intracellular PCA under our current experimental setting **(Fig. S13)**. Third, we found that PCA exhibited notable instability with approximately 20% of degradation regardless of the medium, which considerably contributed to the decreased PCA yield **(Fig. S14)**. Overall, the WY10-pRn showed a rise of 3.33-fold in the titer and 3.90-fold in the yield of PCA than concentrated wild-type cells **(Figs. 3E and S9)**, offering a promising strategy to drive decoupled bioproduction and cell growth with distinct carbon sources derived from waste plant biomass. Due to a natural tolerance up to 15 mM of PCA **(Fig. S15)**, the performance of engineered *R. nubinhibens* may be further improved.

### Cross-module push-and-pull synergistic interactions

To decipher the underlying metabolism, we performed the isotopic labeling analysis with ^13^C-labeled acetate and non-labeled 4HB. The substrates and produced PCA in WY10-pRn with two functional modules (the Experimental group) and in WY10-pCtrl without a bioproduction module (the Control group) were identified from the culture media via mass spectrometry (MS), and the ^13^C-labeled patterns of the intracellular metabolites were characterized by ultra-performance liquid chromatography combined with triple quadrupole mass spectrometry (UPLC-TQ-MS). PCA was completely derived from 4HB with all PCA non-labeled in the reprogrammed WY10-pRn **(Figs. 4A and S16)**, and while a quantitative evaluation cannot be made at current stage, we identified ^13^C-labeled metabolites in the essential carbon metabolic pathways, including the tricarboxylic acid (TCA) cycle, gluconeogenesis, and pentose phosphate (PP) pathway **(Figs. 4B and S17)**, demonstrating the orthogonality between the bioproduction pathway and central metabolism and an efficient metabolic segregation of the two modules. We noticed that 50 - 85% of the carbon originated from ^13^C-labeled acetate, where the non-labeled carbon may be explained by cell growth on the yeast extract. These results were in agreement with the growth profiles of *R. nubinhibens* on acetate and yeast extract, where yeast extract can support cell growth to an approximate OD of 0.3 **(Figs. 2D and 3B).** This was further demonstrated by a control experiment with only ^13^C-labeled acetate and yeast extract as substrates, where the main intracellular metabolites showed similar labeling patterns **(Fig. S18)**. Interestingly, we noticed an abnormal ^13^C-labeled pattern of acetyl-CoA without a functional bioproduction module (without the substrate 4HB or an independent *pobA*) **(Figs. S17 and S18)**, which might suggest a shifted equilibrium of the bidirectional reactions of acetyl-CoA in the presence of yeast extract.

**Fig. 4.**
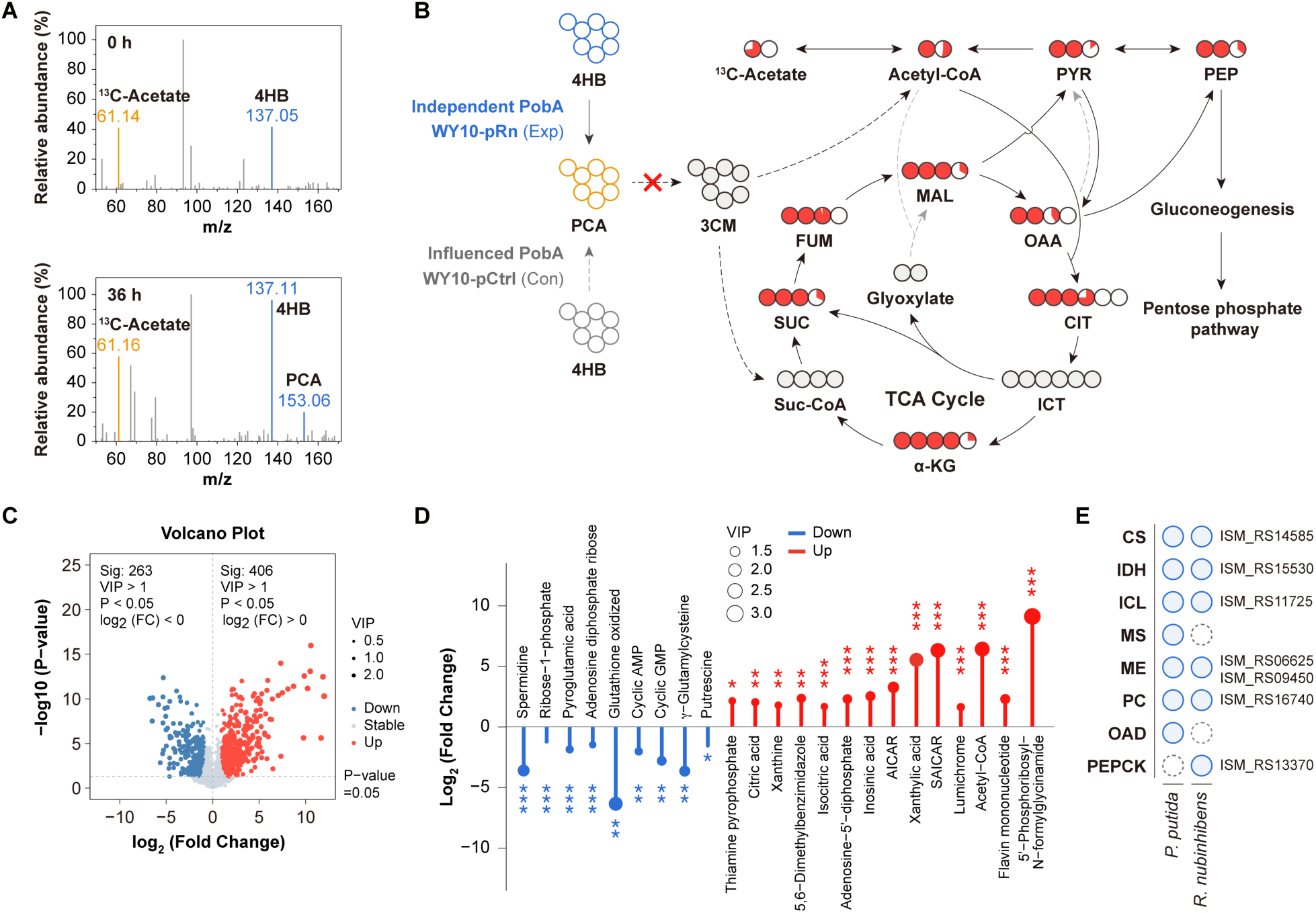
Isotopic labeling analysis and metabolomics of the reprogrammed *R. nubinhibens*. **(A)** MS analysis of the substrates and product of WY10-pRn. The ^13^C-labeled acetate is indicated in purple and the non-labeled 4HB and PCA are indicated in blue. **(B)** Isotopic labeling patterns of metabolites in the central carbon metabolism. The pie chart represents the weighted labeling percent of labeled molecules, indicating the overall contribution of ^13^C-carbon to the carbon source of metabolites. The fractions of non- and ^13^C-labeled carbon are indicated by the white-and purple-filled circles, respectively. The grey-filled circles represent metabolites that were not detected. The grey dashed lines represent catalytical step with genes that have not been identified in the genome of *R. nubinhibens*. Con and Exp represent the control strain WY10-pCtrl and the reprogrammed strain WY10-pRn, respectively. **(C)** Volcano plot. Metabolites with P-value < 0.05 and VIP > 1 were defined as the differential metabolites. The upregulated and downregulated metabolites are indicated by the red and blue dots, respectively. Gray dots represent the metabolites with insignificant changes. **(D)** Lollipop chart of key metabolites with significant changes. The purple and blue bars represent upregulated and downregulated metabolites, respectively, and the size of circles represents the variable importance in projection (VIP) values. *, P<0.05; **, P<0.01; ***, P<0.001. **(E)** Comparison of key genes potentially responsible for metabolic reprogramming in *P. putida* and *R. nubinhibens*. The blue-filled circles represent genes that have been identified and the grey dashed circles represent genes that have not been identified in the genome of *R. nubinhibens*. **Abbreviations:** 3CM, β-carboxy-*cis*-*cis*-muconate; PYR, pyruvate; PEP, phosphoenolpyruvate; MAL, malate; OAA, oxaloacetate; CIT, citrate; ICT, isocitrate; α-KG, α-ketoglutarate; Suc-CoA, succinyl-CoA; SUC, succinate; FUM, fumarate; CS, citrate synthase; IDH, isocitrate dehydrogenase; ICL, isocitrate lyase; MS, malate synthase; ME, malic enzyme; PC, pyruvate carboxylase; OAD, oxaloacetate decarboxylase; PEPCK, phosphoenolpyruvate carboxykinase.

According to the reported acetate metabolism in bacteria, acetate is first converted to acetyl-CoA, and acetyl-CoA is subsequently condensed with oxaloacetate to form citrate, entering the TCA cycle and other pathways, including gluconeogenesis and PP pathway ^46^. Through metabolomics analysis of the intracellular metabolites, 669 metabolites in WY10-pRn showed significant changes compared to WY10-pCtrl, where 406 metabolites were upregulated and 263 metabolites were downregulated **(Figs. 4C and S19)**. While the metabolite pools in the gluconeogenesis and PP pathway remained stable, three metabolites closely related to the TCA cycle, acetyl-CoA, citrate, and isocitrate, were upregulated. Particularly, acetyl-CoA in WY10-pRn elevated 52.08-fold than that in WY10-pCtrl **(Fig. 4D)**. These observations were in agreement with *P. putida* grown on gluconeogenic substrates, where the TCA cycle plays a more important part to fuel cell growth ^47^. These changes also suggest a cofactor-driven metabolic reprogramming, as proposed in a recent report ^19^, to complement the extra consumption of NADPH for the conversion of 4HB to PCA. Indeed, we found the NADP-dependent malic enzyme (encoded by *ISM_RS06625*) and isocitrate dehydrogenase (encoded by *ISM_RS15530*) in the genome of *R. nubinhibens*, which might be responsible for the NADPH generation **(Fig. S20)** ^48^. Interestingly, we observed a lack of oxaloacetate decarboxylase and malate synthase but the existence of phosphoenolpyruvate carboxykinase (encoded by *ISM_RS13370*) in *R. nubinhibens* **(Fig. 4E)**, thereby potentially leading to differently shifted metabolisms. This response to the extra requirement of NADPH was also witnessed by four upregulated metabolites in the riboflavin metabolism, supplying precursors for the flavin adenine dinucleotide as the cofactor of the NADPH-dependent hydroxylation from 4HB to PCA **(Figs. S21 and S22)** ^49–51^. Moreover, seven upregulated and four downregulated metabolites were enriched in the pathway related to purine metabolism, which is essential for nucleic acid synthesis, energy transfer and cofactor-regeneration **(Figs. S21 and S22)**, and one upregulated and five downregulated metabolites were identified in the glutathione metabolism **(Figs. S21 and S22)**, suggesting an integrative adaptation of multiple pathways. Though the exact mechanism of energy and cofactor supplement in the engineered strain remains to be explored, the metabolic response illustrates a cross-module push-and-pull synergy between the genetically orthogonal bioproduction and growth modules, underscoring a unique biological promise for efficient upcycling of waste plant biomass with engineered marine *R. nubinhibens*.

### Dual-modular manipulation enables β-KA production

To extend the utility of our strategy from single enzymatic conversion to pathway-level modulation, we targeted another product β-KA, a precursor for the synthesis of performance-advantaged polymers, by inactivating *pcaIJ* for bioproduction and supplying acetate for cell growth **(Fig. 5A)**. We inactivated these two genes using PAM-independent base editing with dSpRY as the CRISPR-Cas effector rather than dCas9 ^52^, as dCas9 did not work on these loci due to PAM-dependent low efficiency **(Fig. 5B)**. The resulting strain PR71 successfully produced β-KA at 24 h with a yield of 11.27 ± 1.53% with 4HB as substrate and acetate as the carbon source for growth **(Figs. 5C and Table S5)**. As *pcaIJ* is located at a distant locus from the *pca* operon, their inactivation exerted no impact on the pathway upstream. Meanwhile, we overexpressed *pcaHG* independently in PR71 to debottleneck the conversion downstream of PCA **(Figs. 5D and 5E)**. As expected, the *pcaHG*-overexpressing PR71 showed substantially reduced PCA accumulation and the maximal β-KA titer reached to 1.64 ± 0.09 mM **(Fig. 5E)**.

**Fig. 5.**
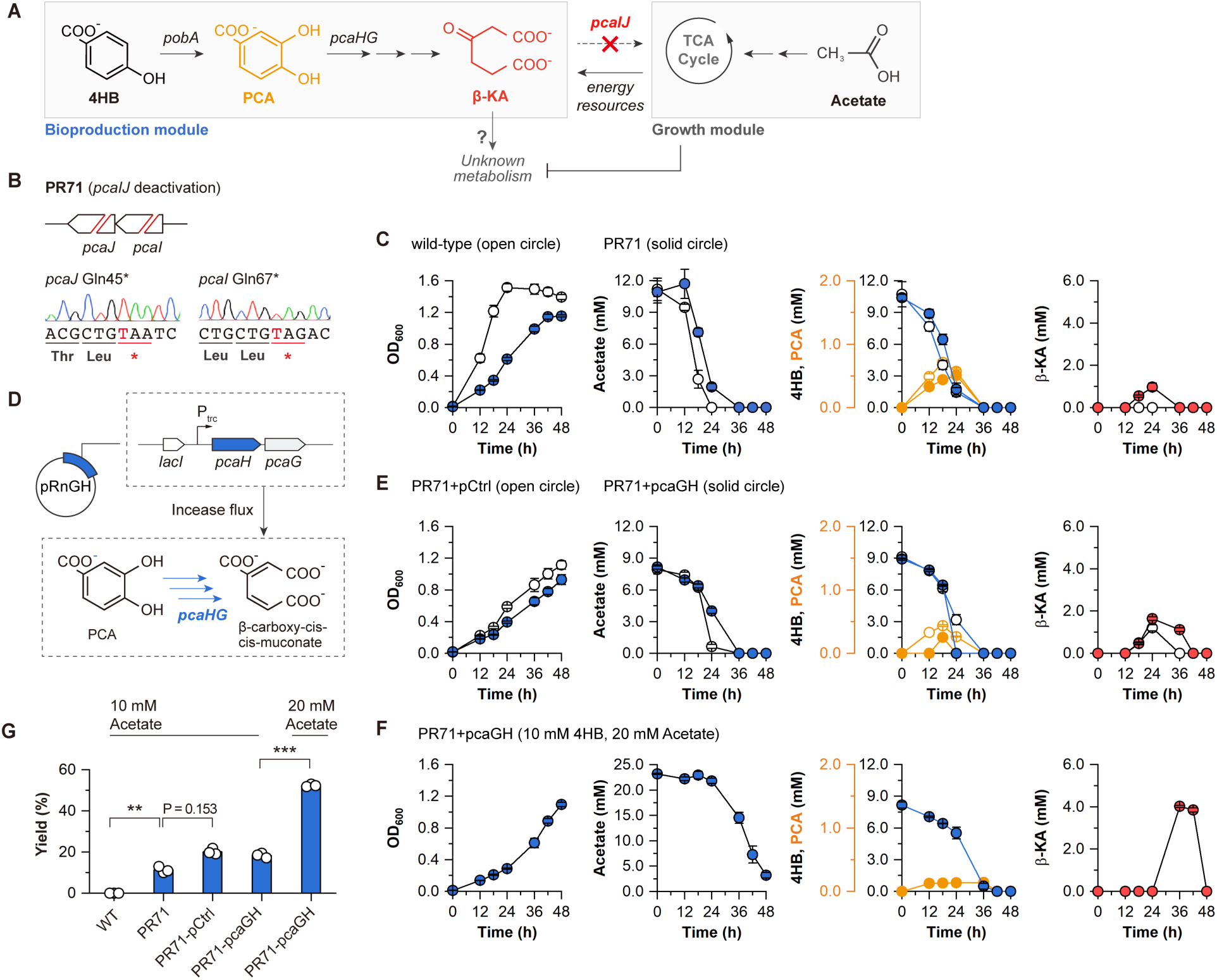
Dual-modularly modulation for β-KA bioproduction. **(A)** Design scheme of the bioproduction and cell growth modules for β-KA production. 4HB was converted to β-KA through the β-ketoadipate pathway with deactivated *pcaIJ* blocking β-KA entering into the TCA cycle, while β-KA may be metabolized through unknown pathways. Acetate is utilized for cell growth. **(B)** Genome editing results of *pcaI* and *pcaJ* in PR71. Based on the wild-type *R. nubinhibens*, the *pcaI* and *pcaJ* genes were deactivated with base editing by introducing early STOP codons in both genes, generating PR71 (*pcaI* Gln67\**, pcaJ* Gln45*). The introduced STOP codons are indicated in red. **(D)** Design of the *pcaHG*-overexpressing plasmid pRnGH. The *pcaG* and *pcaH* genes from *R. nubinhibens* were overexpressed with a plasmid, controlled by an inducible *lacl*-P_trc_ system. Bioproduction performance of the **(C)** PR71 and **(E)** PR71-pcaGH with 10 mM 4HB and 10 mM acetate compared to the wild-type and PR71-pcaGH, respectively. **(F)** Bioproduction performance of the PR71-pcaGH with 10 mM 4HB and 20 mM acetate. **(G)** Molar yields of β-KA from 4HB. Samples were taken at different time intervals. The experiments were conducted in triplicate, and the circles and error bars represent the individual values and standard deviations of three biological replicates, respectively. The differences were statistically evaluated by *t*-test (**, P<0.01; ***, P<0.001, unpaired and two-tailed).

Nevertheless, we found that β-KA was metabolized after 24 h of cultivation even though *pcaIJ* were both inactivated **(Figs. 5C and 5E)**. This was in agreement with a β-KA production work in *P. putida* via repressing *pcaIJ* using CRISPR interference ^53^. A possible route would be the conversion of β-KA to levulinic acid through a non-enzymatic reaction, and then the levulinic acid would go through a catabolic pathway into β-oxidation, generating acetyl-CoA and propionyl-CoA ^54^. As reported, efficient β-KA production in *P. putida* needs inactivation of *lvaE* to prevent the levulinic acid metabolism ^25^. However, we could not identify similar pathways in *R. nubinhibens*, nor could we rule out other unknown metabolic routes contributing to the degradation of β-KA. We found that the majority of β KA was degraded only after acetate was consumed to a certain level (3 – 5 mM) **(Figs. 5C and 5E)**, indicating potential sequential utilization of acetate and β-KA in *pcaIJ*-deficient strains. Therefore, we extended our dual-modular strategy by increasing the carbon source for the growth module. We anticipated that increased acetate would not only supply resource but also modulate β-KA degradation. When the initial acetate was 20 mM, the titer of β-KA significantly increased to 4.02 ± 0.05 mM in the *pcaHG*-overexpressing PR71, and the product was not degraded until 42 h **(Fig. 5F)**, showing a 2.85-fold increase of yield compared to that cultivated with 10 mM acetate **(Fig. 5G)**. These results successfully demonstrated that our dual modular system can efficiently repurpose metabolism for bioproduction, although more efforts are still needed to decipher the biology of a novel chassis with evolutionary advantages yet without an in-depth understanding of their physiology and genetics.

### Valorization of lignin-derived monomers with seawater

We envision a biological scheme with reprogrammed marine *R. nubinhibens* as chassis and seawater as the water source to valorize plant biomass carbon, especially lignin-derived monomers, with minimal reliance on freshwater **(Fig. 6A)**. Different from synthetic media containing defined compositions, such as MBM, the components of actual seawater are highly complex, including dissolved inorganic salts, organic matters and trace elements, and are dramatically fluctuating depending on locations, depths and other natural or anthropogenic conditions. The salinity of seawater is similar to MB2216 but higher than MBM **(Table S7)**. Therefore, it is critical to evaluate our designed system with real seawater. As a proof of principle, we collected seawater in the Yellow Sea near our campus, and prepared seawater basal media (SBM) by adding essential nutrients. WY10-pRn exhibited a promising bioproduction competence with 4HB and acetate as the carbon sources, and the strain converted 5.07 ± 0.39 mM 4HB to PCA with the titer of 3.02 ± 0.46 mM and the yield of 59.47 ± 7.89% **(Fig. 6B)**. The slightly dropped titer and yield in SBM might be attributed to the complex components in seawater. Despite the expected superior and stable PCA production, WY10-pRn with a well functional bioproduction module exhibited better performance and growth than WY06-pRn in SBM **(Fig. S23)**, showing adaptive advantages and robustness when transferred into SBM from MBM.

**Fig. 6.**
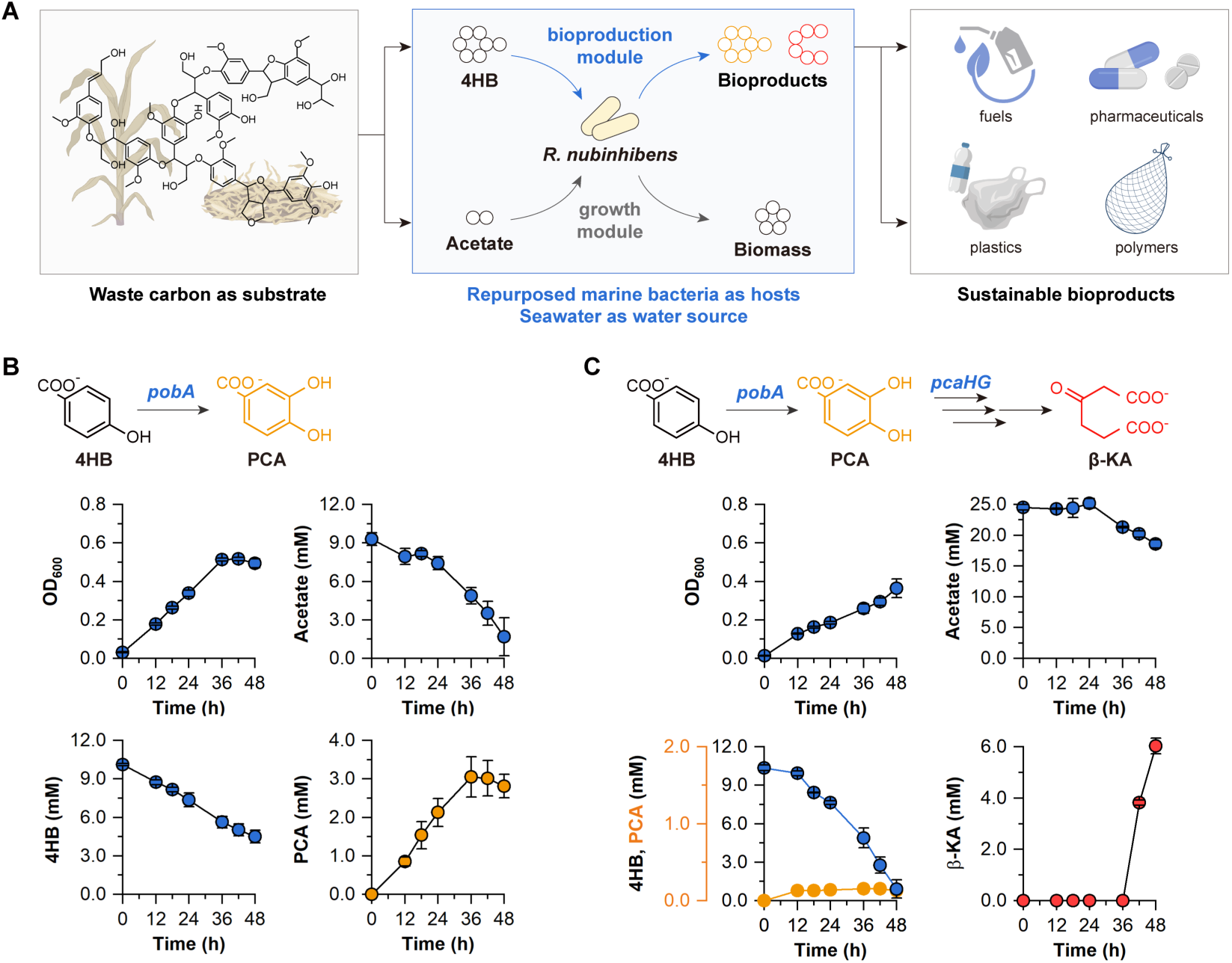
Valorization of lignin-derived monomers with seawater as the water source. **(A)** Schematic illustration of the blue biological scheme to valorize waste plant biomass with modularly reprogrammed *R. nubinhibens*. Bioproduction performance of the **(B)** WY10-pRn and **(C)** PR71-pcaGH with seawater as the water source. The WY10-pRn was cultivated in SBM with 10 mM 4HB and 10 mM acetate. The PR71-pcaGH was cultivated in SBM with 10 mM 4HB and 20 mM acetate. Samples were taken at different time intervals. The experiments were conducted in triplicate, and the circles and error bars represent the individual values and standard deviations of three biological replicates, respectively. Panel **A** created with BioRender.com.

Interestingly, β-KA showed elevated titer and yield in SBM than that in MBM. In PR71 with *pcaHG* overexpression, the titer reached 6.03 ± 0.31 mM and the yield was approaching 64.04 ± 3.27 %, which were 1.50 and 1.22 folds higher than those in MBM **(Figs. 5F and 6C)**. Notably, we did not observe obvious degradation of β-KA within the tested time frame, during which acetate was consumed by 5.97 mM in total and remained above a presumed threshold value of 3 - 5 mM. The difference might also result from the complex components of seawater, which slowed the overall metabolism and enabled elevated bioproduction. We admit that the strain needs further rational engineering and adaptive evolution to reach comparable performance with established hosts ^55,56^, yet the advantages are also appealing in that no freshwater or glucose is necessary at all for this bioproduction. Calculated preliminarily from previous reports ^55,56^, this freshwater- and glucose-independent scheme could save 20 to 40 liters of freshwater and 0.1 to 1.5 kg of glucose for producing 1 kg of β-KA, depending on distinct strategies, whereas seawater and acetate are required instead.

Taken together, these results unraveled the potential of our biosystem using seawater directly as the water source for efficient valorization of lignin-derived monomers. For real-world applications, we envision that acetate and lignin-derived monomers can be obtained from existing processes, such as acid pretreatment and enzymatic conversions. Seawater can be collected and filtered in coastal areas, then supplemented with essential nutrients and the two substrates. The biological conversion, driven by the engineered *R. nubinhibens*, can be performed in a bioreactor operating in either continuous or sequential-batch mode. Given the high-salt nature of the medium, we also foresee open-culture fermentation in the future to further reduce operational costs and energy consumption.

## Discussion

Evolution endows marine microbes with unusual metabolic diversity and environmental adaptability for survival and reproduction in the ocean ^57,58^. The evolutionary fitness, which can hardly be artificially created *de novo* in model microbes, also grants marine microbes great potential for the valorization of diverse yet refractory carbon compounds. For instance, *Alcanivorax* species can degrade alkanes ^59^, and *Roseobacter* clade bacteria can metabolize carbon monoxide, aromatic compounds, and dimethylsulfoniopropionate ^60^. Recent studies report that coastal microbial consortia can naturally degrade lignocellulosic biomass, demonstrating their inherent potential for waste carbon degradation ^61,62^. These advantages are particularly compelling under conditions mimicking the ocean, establishing the foundation for water-conserving bioprocesses. By leveraging the evolutionary fitness, we establish in the present study a blue biological scheme for the integrative conservation of carbon and water, enabling the efficient valorization of lignin-derived carbon with seawater, rather than freshwater, as the water source.

However, the evolutionary fitness also accompanies unexpected challenges preventing the establishment of engineered functionalities. We discovered a rare polar effect in *R. nubinhibens*, in which the deactivation of any genes in the *pca* cluster would interfere with others in the same operon. The superior catalytic property of the PobA from *R. nubinhibens* can only be released by circumventing the polar effects. Therefore, we reprogrammed the independency of *pobA* and genetically decoupled metabolic network, achieving a 13.17-fold increase in the yield of PCA (65.85%) over natural accumulation of the wild type (5.00%) ^12^. This was achieved by utilizing a broad-host, low-copy number plasmid, pBBR1MCS-5, which is currently the only available and stable vector series for the *Roseobacter* clade bacteria, with tunable expression driven by the *lacI*-P_trc_ inducible system. Future endeavors require more accessible genetic systems (e.g., plasmids with different copy numbers, regulatory parts, and gene editing tools) to engineer the intrinsic features of bacteria. Indeed, we reported previously the failure of CRISPR-Cas counterselection for gene insertions and deletions, indicating unanticipated difficulties in developing genetic systems in bacteria with unusual evolutionary features. Instead, the successful utilization of deamination-mediated base editing for single-nucleotide resolution modifications in *R. nubinhibens* and other strains suggests a practicable path for engineering marine bacteria or other non-model strains ^37^, enabling maximal designed outcomes with minimal genetic changes.

Due to the metabolic diversity, marine microbes tend to channel all accessible resources to cell growth rather than bioproduction of chemicals for human needs. Metabolically reprogramming carbon allocation often leads to transient accumulation of products or failure in both bioproduction and cell growth with a single carbon source. This was observed when we previously produced PCA using a strain with CRISPRi-regulated metabolism ^12^, and when in this study we attempted to produce β-KA in PR71 with both subunits of the downstream enzyme being inactivated. The desired products were eventually metabolized in both cases. Furthermore, most metabolic pathways cannot be thoroughly independent to cell growth due to the requirement of enzymes, cofactors and energy, while it is possible to direct the primary carbon source to a designed route with a second carbon source sustaining cell growth ^63^. As such, we generated orthogonal bioproduction and growth modules in *R. nubinhibens* via CRISPR-Cas base editing at the single-nucleotide resolution and introduced acetate to intentionally sustain cell growth and provide necessary resources for bioproduction, which was solely driven by lignin-derived carbon. While the orthogonality of bioproduction and cell growth was demonstrated via ^13^C labeling analysis, we observed cross-module push-and-pull synergistic interactions between these two modules besides the designed metabolic segregation. This is not surprising that the growth module guarantees adequate resources pushing bioproduction, in which the resource consumption, in return, pulls the central metabolism. By doing so, the bioproduction of PCA was significantly elevated by 8.28-fold in titer than that of the wild-type strain (0.46 mM) ^12^. Interestingly, we report for the first time that supplying a second carbon source for growth can effectively delay the degradation of intended bioproducts without disrupting the unknown metabolic route. The β-KA titer and yield increased by 2.46- and 2.85-folds **(Figs. 5F and 5G)**, respectively, when supplying 20 mM acetate, and the performance was even more appealing with seawater as a medium, reaching 6.03 ± 0.31 mM and 64.04 ± 3.27 % **(Fig. 6C)**. While a systematic interrogation of possible metabolic routes is necessary to achieve higher and more stable bioproduction, our strategy provides a promising alternative by continuously supplying substrates to the growth module. This would be particularly beneficial for non-model hosts with evolutionary advantages yet without an in-depth deciphering of their metabolism.

Repurposing waste carbon into valuable products with engineered microbial chassis holds a grand promise to establish a circular bioeconomy. This is particularly crucial for the valorization of waste plant biomass, of which traditional disposals (e.g., via incineration) not only aggravate carbon emissions but also exert severe environmental issues. With our system, one primary yet recalcitrant aromatic carbon and a universal byproduct from waste plant biomass can be valorized to pharmaceuticals without requesting immediately fermentable carbon nor reliance on freshwater. By utilizing seawater as the water source, we can substantially reduce freshwater reliance in next-generation biotechnology, providing an integrative biological scheme leveraging precisely reprogrammed microbes with unusual competencies for achieving carbon and water conservation goals simultaneously. Looking beyond our immediate goals, we envision that this blue biological scheme could enable the implementation of next-generation biotechnology in regions where basic water supply for human needs is scarce.

## Methods

### Strains and media

All strains used in this study were summarized in **Table S5**. *E. coli* DH5α (Takara) was used for molecular cloning and *E. coli* BW25113 was used for expressing different *pobA* genes. *P. putida* KT2440 was used for comparison with *R. nubinhibens*. Luria-Bertani (LB) media were used to cultivate *E. coli* at 37 °C and 180 rpm. *P. putida* was grown in LB for routine cultivation, and in SBM for performance comparison, at 30 °C and 180 rpm. *R. nubinhibens* ISM was grown in MB2216 for routine cultivation, and in MBM and SBM for bioproduction, at 30 °C and 180 rpm. Detailed compositions of the above-mentioned media were as described previously ^12^. Ampicillin (100 μg/mL) or gentamicin (20 μg/mL) was supplemented for transformant selection or plasmid maintenance when necessary.

### Plasmid and strain construction

All plasmids, primers, synthesized gene sequences and gRNAs sequences, used in this study were summarized in **Table S2**, **Table S8**, **Table S9** and **Table S10**, respectively. The *pobA* genes from *P. putida* and *C. glutamicum* were commercially synthesized with their original codons using pUC57 as the backbone plasmid. The *pobA*, *pcaH* and *pcaG* genes from *R. nubinhibens* were amplified from its genome. To construct pPp, pCg and pRn, three *pobA* fragments were individually fused into the pCtrl backbone carrying the *lacI*-P_trc_ inducible system. By replacing *pobA* with *pcaH* (or *pcaG*), the plasmid pRnH (or pRnG) was generated. By replacing *pobA* with *pcaG* and *pcaH*, the plasmid pRnGH was generated. All DNA fragments were amplified with PrimeSTAR Max DNA Polymerase (Takara) and fused with In-Fusion Snap Assembly Master Mix (Takara). All plasmids were extracted with QIAprep Spin Miniprep Kit (Qiagen).

Electroporation was used to deliver plasmids into *R. nubinhibens* as described previously ^37^. All base editing plasmids carried the *lacI*-P_trc_ inducible system, PmCDA1 deaminase and uracil glycosylase inhibitor, together with either dCas9 or dSpRY, and a customized gRNA cassette targeting the gene of interest. To generate WY10 (*pcaH* Gln40*, *pcaG Gln3\**), the previously constructed plasmid pWY07 ^12^, carrying a dCas9-based editor and a customized gRNA (gRNA-pcaG) targeting *pcaG*, was transformed into WY06 (*pcaH* Gln40*). To generate dSpRY-based editor, dCas9 in plasmid pWY was replaced with dSpRY ^52^, generating the template plasmid pdSpRY. Customized gRNA cassettes were subsequently integrated into pdSpRY to construct the editing plasmids pRC42 (with gRNA-pcaI) and pRC43 (with gRNA-pcaJ), targeting *pcaI* and *pcaJ*, respectively. The plasmid pRC42 was transformed into wild-type *R. nubinhibens*, generating PR70 (*pcaI* Gln67*), and the plasmid pRC43 was transformed into PR70 (*pcaI* Gln67*), generating PR71 (*pcaI Gln67*,* pcaJ *Gln45\**). All base editing was induced with 0.5 mM IPTG, and the successful introduction of early STOP codons (TAA or TAG) was confirmed by Sanger sequencing the targeted loci. All gRNA cassettes were constructed by inverse PCR using back-to-back primers containing 20 bp spacer sequences. For plasmid curing, the edited strain was cultivated in the MB2216 liquid media without gentamicin and streaked on the MB2216 solid media. The successful plasmid curing was determined by the loss of signal of the working plasmid and the recovered sensitivity to gentamicin.

### Phylogenetic analysis

The PobA proteins of the lignin-valorizing microbes and representative bacteria from the marine *Roseobacter* clade were retrieved from the National Center for Biotechnology Information, and nine PobA amino acid sequences were chosen for phylogenetic analysis **(Table S11)**. Sequence alignment was performed with ClustalW and a maximum likelihood phylogenetic tree was built with 1,000 bootstrap replicates using MEGA11.

### Evaluation of enzyme activity

The strains carrying different *pobA* genes were cultivated in MB2216 overnight and were harvested and washed twice with precooled phosphate buffered saline (PBS) or MBM-Tris (Tris was dissolved in MBM). The resuspended cells were disrupted via ultrasonication on ice. After removing cell debris by centrifugation at 15,000 rpm for 10 min at 4 °C, the supernatant, as the crude cell extract, was stored at -80 °C until protein and enzyme activity assay. The total protein was determined using BCA Protein Assay Kit (Solarbio). The enzyme activity of PobA was determined by the oxidation of NADPH or NADH based on its absorbance at 340 nm. For the low-salinity reaction system, the reaction mixture contained 75 mM Tris, 0.5 mM NADPH or NADH, 5 mM MgCl_2_, 10 mM 4HB and 20 μL of cell extract. For the high-salinity reaction system, Tris was replaced with MBM-Tris, and MgCl_2_ was removed. The pH of the reaction system was adjusted with HCl according to the experimental setting. The reaction was incubated at 30 °C for 2 h and the absorbance at 340 nm was measured with intervals of 20 s by Spark Multimode Microplate Reader (Tecan). One unit of the PobA enzyme activity was defined as the amount of enzyme catalyzing the reaction of 1 μM substrate per minute.

### Molecular docking

Molecular docking was performed to investigate the interactions between the protein (PobA) and substrate (4HB) using AutoDock 4.2 software. The pristine sequences and corresponding PDB files were collected from AlphaFold under accession numbers Q88H28 (PpPobA), A0A1Q6BNG9 (CgPobA) and Q6SJC7 (RnPobA). Subsequently, all these files are submitted to H++ 4.0 package (http://biophysics.cs.vt.edu/H++) to correct the structures based on the predicted pKa values of the amino acid residues. The grid-point and grid maps were set to 0.375 Å and 180 × 180 × 180, respectively. The population size, maximum number of evaluations, and maximum number of generations were set to 150, 2,500,000, and 27,000 with the other default parameters automatically provided by the software. As for each docking case, 100 independent runs were conducted to identify the most stable configuration with the lowest binding free energy using Lamarckian genetic algorithm. The final protein-ligand configurations were visualized by PyMOL.

### Quantification of the transcription level of *pobA*

All primers used in this study were summarized in **Table S8**. The wild-type and *pcaHG*-deficient *R. nubinhibens* were cultivated in MB2216 with 4HB for 12 h, and cells were collected at the exponential phase with the OD_600_ at 0.5. Total RNA was extracted with RNAprep Pure Cell/Bacteria Kit (Tiangen), and then was reverse-transcribed to cDNA for analysis with the PrimeScript RT reagent Kit with gDNA Eraser (Perfect Real Time) (Takara Bio). Quantitative PCR was conducted with TB Green Premix Ex Taq II (Tli RNaseH Plus) (Takara Bio) using Applied Biosystems QuantStudio 5 (Thermo Fisher Scientific). Gene expression levels across samples were normalized using the 16S rRNA gene as the housekeeping gene.

### Cultivation and bioproduction experiment

The control and experimental *R. nubinhibens* strains were first cultivated in MB2216 overnight before subsequent experiments. For performance comparison of *P. putida* and *R. nubinhibens*, wild-type strains were cultured in 50 mL SBM with 10 mM 4HB. The pH of SBM was adjusted to 8.1 or 8.5 with NaOH according to the experimental setting. For cell growth of *R. nubinhibens* on sugar substrates, the wild-type strain was cultured in 30 mL MBM with 5 g/L glucose, 5 g/L xylose, 10 mM 4HB or without additional carbon source. For PCA tolerance of *R. nubinhibens*, the wild-type strain was cultured in 30 mL MB2216 with PCA at various concentrations (0, 5, 10, 15, and 20 mM). For simultaneous bioproduction and cell growth, strains were inoculated into 50 mL fresh MBM with a dilution rate of 1:50 with 10 mM 4HB as the sole carbon source. For two-step bioproduction and cell growth, strains were inoculated into 50 mL fresh MB2216 with a dilution rate of 1:50. When the cells reached the late exponential phase to an OD_600_ of 1.0, the cells were harvested and washed twice with PBS and resuspended in 10 mL MBM with 10 mM 4HB. For the two-carbon utilizing bioproduction and cell growth, strains were inoculated into 50 mL fresh MBM or SBM with a dilution rate of 1:50 with 10 mM 4HB as the substrate for bioproduction and 10 mM (or 20 mM) acetate for cell growth. The pH was adjusted to 7.5 with NaOH. The gentamicin (20 μg/mL) and IPTG (0.5 mM) were added to maintain plasmids and induce the expression of *pobA* when needed.

During cultivation, samples were collected at different time intervals, and subsequently determined for growth profiles via OD_600_ and for the extracellular substrates and products via high performance liquid chromatography (HPLC). After collecting the samples, the OD_600_ of all samples was first measured. Then, the samples were centrifuged at 12,000 rpm for 1 min and the supernatant was collected via filtration (0.22-μm). The OD_600_ of the supernatant was measured again and the real OD_600_ of samples was calculated by deducting the OD_600_ of the supernatant from that of the unprocessed sample. The yield of PCA from 4HB was calculated at the maximal value for each strain. Three independent biological replicates were carried out and the mean, individual values and standard deviations were reported.

### Quantification of 4HB, PCA, acetate and β-KA

4HB, PCA, and acetate were quantified with HPLC (Agilent Technologies). For quantification of the extracellular compounds, the samples were first centrifuged at 12,000 rpm for 1 min and the supernatant was filtered with 0.22 μm filters. For quantification of the intracellular compounds, the cells were washed twice and resuspended with PBS. Next, the cells were disrupted via ultrasonication on ice, and after removing cell debris, the supernatant was stored at -20 °C until analysis. 4HB and PCA were measured at 210 nm by a variable wavelength detector with EC-C18 column (4.6 × 100 mm, 4 μm, Agilent Technologies). The mobile phase consisted of 10% acetonitrile and 90% formic acid (0.1%), and the flow rate was 0.8 mL/min. For PCA production, acetate was measured alone by a refractive index detector (RID) with Aminex HPX-87P column (7.8 × 300 mm, 9 μm, Bio-Rad) at 50 °C. To quantification β-KA, samples were first converted to levulinic acid by adding 10 μL of 5 N H_2_SO_4_ per 0.5 mL of sample, followed by incubation at 50 °C for 8 h. β-KA acid concentration is calculated based on the quantified levulinic acid, assuming a 1:1 molar conversion ^56^. Acetate and levulinic acid were then measured together by a RID with the same Aminex HPX-87P column at the optimized temperature of 30 °C for better separation **(Figure S24).** The mobile phase consisted of 5 mM sulfuric acid and the flow rate was 0.6 mL/min.

### Stability evaluation of PCA

The PCA stability was evaluated in 5 mL different media, including the rich medium MB2216, the synthetic minimal medium MBM, the seawater-based minimal medium SBM and after-filtration seawater alone. PCA was added to all these media to the final concentration at 5 mM and its concentration was monitored via HPLC every 12 h.

### Characterization of ^13^C-labeled metabolites

The non-labeled 4HB and ^13^C-labeled acetate (Sigma-Aldrich, Product No. 282014, CAS No. 56374-56-2) were added to MBM to cultivate WY10-pCtrl for 20 h and WY10-pRn for 36 h to reach mid exponential phase with the OD_600_ at 0.5 for isotopic labeling analysis of the intracellular metabolites. To confirm that the non-labeled fractions were derived from the yeast extract and not from 4HB, only ^13^C-labeled acetate without 4HB was added to MBM to cultivate WY10-pRn for 12 h at the mid exponential phase with the OD_600_ at 0.5. The ^13^C-labeled patterns of 4HB and PCA in the culture media were characterized by MS (Ultimate 3000, ISQ EC, Thermo Fisher Scientific). Before MS analysis of the extracellular metabolites, samples were first centrifuged at 12,000 rpm for 1 min and the supernatant was filtered with 0.22 μm filters and demineralized with solid phase extraction using HyperSep C18 (Thermo Fisher Scientific). The ^13^C-labeled patterns of key metabolites involved in the central carbon metabolism were characterized by UPLC-TQ-MS (ACQUITY UPLC I-Class/Xevo TQ-S, Waters) with XBridge BEH HILIC column (2.1 × 100 mm, 2.5 μm, Waters). The cells were harvested and washed twice with precooled PBS at 4 °C and were quickly frozen with liquid nitrogen. Then, cells were resuspended with 80% methanol and were disrupted via ultrasonication. After removing cell debris by centrifugation, the supernatant was concentrated and resuspended with 80% methanol for UPLC-TQ-MS analysis.

### Metabolomics analysis

The WY10-pCtrl and WY10-pRn strains were cultivated in MBM with 4HB and acetate for 20 h and 36 h with the OD_600_ at 0.5, respectively, for metabolomics analysis of the intracellular metabolites. To prepare the samples, cells were harvested and washed twice with precooled PBS at 4 °C and were quickly frozen with liquid nitrogen. The metabolites were analyzed by LC-MS (ACQUITY UPLC I-Class PLUS, Waters/QE, Thermo) with an ACQUITY UPLC HSS T3 column (2.1 × 100 mm, 1.8 μm, Waters). Data were processed using Progenesis QI v3.0, and metabolites were identified based on The Human Metabolome Database, Lipidmaps (v2.3), METLIN and LuMet-Animal databases. Metabolites with P-value < 0.05 and VIP > 1 were defined as the differential metabolites. Pathway enrichment analysis of differential metabolites was performed based on KEGG database.

## Data availability

The metabolomics data are available at MetaboLights repository under accession number MTBLS12392.

## Acknowledgments

This work was supported by the National Natural Science Foundation of China (22578250, 22278246 and 22378233), the Intramural Joint Program Fund of State Key Laboratory of Microbial Technology (Project NO. SKLMTIJP-2025-02 and SKLMTIJP-2024-01), and the Taishan Scholars Project of Shandong Province (NO. tstp20230604).

