## Supporting information for "Repurposing the evolutionary fitness of marine bacteria for water-conserving conversion of lignin-derived carbon"

**Table S1.** The percent identities of genes from *R. nubinhibens* and *P. putida*.

| | **Expect value** | **Percent identities** |
|---|---|---|
| *pobA* | 8e-50 | 65.92% |
| *pcaH* | 4e-55 | 67.29% |
| *pcaG* | 2e-13 | 80.95% |

**Table S2.** Plasmids used in this study.

| Name | Description | Source |
|---|---|---|
| pUC57-Pp | *ColE1* ori, *bla*, *pobA-Pp* | Beijing Tsingke Biotech Co., Ltd. |
| pUC57-Cg | *ColE1* ori, *bla*, *pobA-Cg* | Beijing Tsingke Biotech Co., Ltd. |
| pBBR1MCS-5 (pCtrl) | pBBR1 ori, pBBR1 Rep, GmR | Lab stock |
| pTemplate | pUC ori, *bla*, gRNA scaffold | Lab stock |
| pmCherry | pCtrl, *lacI*-$P_{trc}$, mCherry | Lab stock |
| pWY | pCtrl, *lacI*-$P_{trc}$, *dcas9*, *PmCDA1*, *ugi* | Lab stock |
| pWY07 | pWY, gRNA-pcaG | Lab stock |
| pdSpRY | pCtrl, *lacI*-$P_{trc}$, *dSpRY*, *PmCDA1*, *ugi* | This study |
| pgRNA-pcaI | pTemplate, gRNA-pcaI | This study |
| pgRNA-pcaJ | pTemplate, gRNA-pcaJ | This study |
| pRC42 | pdSpRY, gRNA-pcaI | This study |
| pRC43 | pdSpRY, gRNA-pcaJ | This study |
| pPp | pCtrl, *lacI*-$P_{trc}$, *pobA* from *P. putida* | This study |
| pCg | pCtrl, *lacI*-$P_{trc}$, *pobA* from *C. glutamicum* | This study |
| pRn | pCtrl, *lacI*-$P_{trc}$, *pobA* | This study |
| pRnH | pCtrl, *lacI*-$P_{trc}$, *pcaH* | This study |
| pRnG | pCtrl, *lacI*-$P_{trc}$, *pcaG* | This study |
| pRnGH | pCtrl, *lacI*-$P_{trc}$, *pcaH*, *pcaG* | This study |

**Table S3.** The composition of Tris and MBM-Tris buffer.

| | **Tris** | **MBM-Tris** | **Seawater** |
|---|---|---|---|
| Tris | 75 mM (9.09 g/L) | 75 mM (9.09 g/L) | |
| NaCl | / | 160 mM (9.35 g/L) | |
| $MgSO_4$ | / | 40 mM (4.82 g/L) | |
| $MgCl_2$ | 5 mM (0.48 g/L) | / | |
| $CaCl_2$ | / | 8 mM (0.89 g/L) | 35 g/L |
| KCl | / | 8 mM (0.6 g/L) | |
| $NH_4Cl$ | / | 8 mM (0.42 g/L) | |
| $K_2HPO_4$ | / | 0.8 mM (0.14 g/L) | |

**Table S4.** Proton numbers and sites of the PobA at different pHs.

| | pH | Proton number | Proton site |
|---|---|---|---|
| PpPobA | Initial | 6232 | None |
| | 7.5 | 6234 | His130 and His280 |
| | 8.0 | 6232 | None |
| | 8.5 | 6232 | None |
| | 9.0 | 6232 | None |
| CgPobA | Initial | 6162 | None |
| | 7.5 | 6163 | His245 |
| | 8.0 | 6163 | His245 |
| | 8.5 | 6163 | His245 |
| | 9.0 | 6163 | His245 |
| RnPobA | Initial | 6018 | None |
| | 7.5 | 6022 | His89, His125, His129 and His288 |
| | 8.0 | 6020 | His89 and His288 |
| | 8.5 | 6019 | His89 |
| | 9.0 | 6018 | None |

**Table S5.** Strains used in this study.

| Name | Feature | Source |
|---|---|---|
| *E. coli* DH5$\alpha$ | Commercial strain for molecular cloning | Takara Bio (Dalian) Co., Ltd. |
| *E. coli* BW25113 | *E. coli* K-12 derivative | Lab stock |
| *P. putida* KT2440 | *P. putida* mt-2 without pWW0 plasmid | Lab stock |
| *R. nubinhibens* ISM | Wild type | Lab stock |
| WY06 | *R. nubinhibens* ISM, *pcaH* Gln40* | Lab stock |
| WY07 | *R. nubinhibens* ISM, *pcaG* Gln3* | Lab stock |
| WY10 | *R. nubinhibens* ISM, *pcaH* Gln40*, *pcaG* Gln3* | This study |
| PR70 | *R. nubinhibens* ISM, *pcaI* Gln67* | This study |
| PR71 | *R. nubinhibens* ISM, *pcaI* Gln67*, *pcaJ* Gln45* | This study |

**Table S6.** Biosynthetic performance of *R. nubinhibens*. [a]

| Strain | Pattern of biosynthesis and cell growth | Initial $OD_{600}$ | Titer (mM) | Yield (%, moL/moL) | Productivity (mM/h) |
|---|---|---|---|---|---|
| WT | Two steps | 5.0 | 1.15 ± 0.16 | 16.97 ± 3.24 | 0.29 ± 0.04 |
| WY06 | Two steps | 5.0 | 0 | 0 | 0 |
| WY06-pCtrl | / | 0.1 | 0 | 0 | 0 |
| WY06-pRn | / | 0.1 | 1.26 ± 0.02 | 77.32 ± 8.23 | 0.06 ± 0.00 |
| WY06-pCtrl | Two steps | 5.0 | 0 | 0 | 0 |
| WY06-pRn | Two steps | 5.0 | 2.91 ± 0.03 | 71.96 ± 9.40 | 1.46 ± 0.02 |
| WY06-pCtrl | Carbon co-utilization | 0.1 | 0 | 0 | 0 |
| WY06-pRn | Carbon co-utilization | 0.1 | 2.77 ± 0.14 | 62.25 ± 5.21 | 0.07 ± 0.00 |
| WY10-pRn | Carbon co-utilization | 0.1 | 3.81 ± 0.43 | 65.85 ± 5.47 | 0.08 ± 0.01 |

[a] Data are the averages and standard deviations from three independent biological experiments.

**Table S7.** The main composition of MB2216, MBM, SBM and Seawater.

<table>
<tr><th></th><th>MB2216</th><th>MBM</th><th>SBM</th><th>Seawater</th></tr>
<tr><td>Peptone</td><td>5 g/L</td><td>/</td><td>/</td><td>/</td></tr>
<tr><td>Yeast extract</td><td>1 g/L</td><td>0.5 g/L</td><td>0.5 g/L</td><td>/</td></tr>
<tr><td>NaCl</td><td>19.45 g/L</td><td>11.69 g/L</td><td rowspan="8">35 g/L</td><td rowspan="8">35 g/L</td></tr>
<tr><td>$MgSO_4$</td><td>3.24 g/L</td><td>6.02 g/L</td></tr>
<tr><td>$MgCl_2$</td><td>5.9 g/L</td><td>/</td></tr>
<tr><td>$CaCl_2$</td><td>1.8 g/L</td><td>1.11 g/L</td></tr>
<tr><td>KCl</td><td>0.55 g/L</td><td>0.75 g/L</td></tr>
<tr><td>$NH_4Cl$</td><td>/</td><td>0.53 g/L</td></tr>
<tr><td>$K_2HPO_4$</td><td>/</td><td>0.17 g/L</td></tr>
<tr><td>$NaHCO_3$</td><td>0.16 g/L</td><td>/</td></tr>
</table>

**Table S8.** Primers used in this study.

| Primer | Sequence |
|---|---|
| **Primers for plasmid construction** | |
| XIA-WY-242 | TTTCACACAGGAAACAGACCATGAAAACTCAGGTTGCA |
| XIA-WY-243 | CGCTTACAATTTCCATTCGCTCAGGCAACTTCCTCGAA |
| XIA-WY-244 | TGCAACCTGAGTTTTCATGGTCTGTTTCCTGTGTGAAA |
| XIA-WY-245 | TTCGAGGAAGTTGCCTGAGCGAATGGAAATTGTAAGCG |
| XIA-WY-246 | TTTCACACAGGAAACAGACCATGAACCACGTACCAGTG |
| XIA-WY-247 | CGCTTACAATTTCCATTCGCTTATACCTCGAAGCGTGG |
| XIA-WY-248 | CACTGGTACGTGGTTCATGGTCTGTTTCCTGTGTGAAA |
| XIA-WY-249 | CCACGCTTCGAGGTATAAGCGAATGGAAATTGTAAGCG |
| XIA-WY-289 | TTTCACACAGGAAACAGACCATGAAAACGCCCGCCGAATA |
| XIA-WY-290 | CGCTTACAATTTCCATTCGCTCAGCCTCCCTCGGGGCGGTTTT |
| XIA-WY-291 | TATTCGGCGGGCGTTTTCATGGTCTGTTTCCTGTGTGAAA |
| XIA-WY-292 | AAAACCGCCCCGAGGGAGGCTGAGCGAATGGAAATTGTAAGCG |
| XIA-WY-293 | TTTCACACAGGAAACAGACCATGGCGCAACGTCTCGACTA |
| XIA-WY-294 | CGCTTACAATTTCCATTCGCCTAGATGTCGAAAAACACGG |
| XIA-WY-295 | TAGTCGAGACGTTGCGCCATGGTCTGTTTCCTGTGTGAAA |
| XIA-WY-296 | CCGTGTTTTTCGACATCTAGGCGAATGGAAATTGTAAGCG |
| XIA-PRC-454 | GCTGCAGACGCGGCAGGTGAGTTTTAGAGCTAGAAATAGC |
| XIA-PRC-455 | TCACCTGCCGCGTCTGCAGCGCTAGCATTATACCTAGGAC |
| XIA-PRC-458 | CAATCAGAAAACGGGATGCTGTTTTAGAGCTAGAAATAGC |
| XIA-PRC-459 | AGCATCCCGTTTTCTGATTGGCTAGCATTATACCTAGGAC |
| XIA-PRC-460 | AGCGGATTTGAACGTTGCGAATCCTTGACAGCTAGCTCAG |
| XIA-PRC-461 | CTGAGCTAGCTGTCAAGGATTCGCAACGTTCAAATCCGCT |
| XIA-PRC-462 | AAGCACACGGTCACAGGAAACAGCTATGACCGTCTCGGTT |
| XIA-PRC-463 | AACCGAGACGGTCATAGCTGTTTCCTGTGACCGTGTGCTT |
| XIA-PRC-493 | TTTCACACAGGAAACAGACCATGAAAACGCCCGCCGAATA |
| XIA-PRC-494 | CGCTTACAATTTCCATTCGCCTAGATGTCGAAAAACACGG |
| XIA-PRC-495 | TATTCGGCGGGCGTTTTCATGGTCTGTTTCCTGTGTGAAA |
| XIA-PRC-496 | CCGTGTTTTTCGACATCTAGGCGAATGGAAATTGTAAGCG |

| | |
|---|---|
| **Primers for colony PCR** | |
| XIA-WY-298 | CGGCTCATCACCCAATGTTA |
| XIA-WY-299 | GGCATCAACATCGGTCTCAA |
| XIA-WY-300 | AAGTGCGCTGTTCCAGACTA |
| XIA-PRC-466 | GTCACAACCCCTCCATCTTA |
| XIA-PRC-467 | GGTGCAGGATTTGAGCAGTT |
| XIA-PRC-468 | TTGCGGCATTCCCGAACTTC |
| XIA-PRC-469 | GCATCGAGCAACGCACTGTA |
| **Primers for qPCR** | |
| XIA-WY-344 | CCTGATCTAGCCATGCCG |
| XIA-WY-345 | CGTATTACCGCGGCTGCT |
| XIA-WY-346 | ATCGACTGCGATTTCATCGC |
| XIA-WY-347 | TAGATCAGCTCGTCATGGAC |

**Table S9.** Sequences of synthesized *pobA* genes used in this study.

| Gene | Sequence [a] |
|---|---|
| *pobA* [b] | ATGAAAACTCAGGTTGCAATTATTGGTGCAGGTCCGTCTGGCCTGCTGCT<br>GGGCCAGCTGCTGCACAAGGCCGGTATCGATAACATCATCGTCGAACGC<br>CAGACTGCCGAGTACGTACTAGGCCGCATCCGCGCCGGGGTGCTAGAG<br>CAAGGCACGGTCGACCTGCTGCGCGAGGCTGGCGTGGCCGAGCGCATG<br>GACCGTGAAGGCCTGGTGCACGAGGGGGTTGAACTGCTGGTTGGCGGG<br>CGCCGCCAGCGTCTGGATCTCAAAGCCCTGACCGGCGGCAAGACGGTG<br>ATGGTCTACGGCCAGACCGAAGTCACCCGTGACCTGATGCAGGCCCGC<br>GAAGCCAGTGGTGCGCCGATCATTTATTCAGCCGCCAACGTTCAGCCGC<br>ATGAATTGAAAGGCGAGAAGCCCTACCTGACGTTCGAAAAGGATGGCCG<br>GGTGCAGCGGATTGACTGCGACTATATCGCCGGCTGCGACGGCTTCCAC<br>GGTATCTCGCGGCAGAGCATCCCGGAGGGCGTGCTGAAACAGTATGAG<br>CGGGTTTACCCGTTTGGCTGGCTGGGCCTGCTGTCGGACACACCGCCA<br>GTCAATCACGAGTTGATCTACGCCCACCATGAGCGCGGTTTCGCGTTGT<br>GTAGCCAACGCTCGCAAACACGCAGCCGCTACTACCTGCAGGTACCTTT<br>GCAGGATCGGGTCGAGGAGTGGTCTGACGAGCGTTTCTGGGACGAACT<br>GAAAGCCCGTCTGCCCGCCGAGGTGGCGGCGGACCTGGTCACAGGCCC<br>CGCGTTGGAAAAAAGTATTGCGCCGCTGCGTAGCCTGGTGGTCGAACCC<br>ATGCAGTATGGTCACCTGTTCCTGGTGGGGGACGCGGCGCACATCGTCC<br>CCCCTACGGGTGCCAAAGGCCTTAACCTGGCGGCCTCCGACGTCAACTA<br>CCTGTACCGCATTCTGGTCAAGGTGTACCACGAAGGGCGCGTCGACCTG<br>CTTGCGCAATACTCGCCGCTGGCACTGCGCCGCGTGTGGAAGGGCGAG<br>CGCTTCAGCTGGTTCATGACCCAACTGCTGCATGACTTCGGTAGCCACAA<br>GGACGCCTGGGACCAGAAGATGCAGGAAGCTGACCGCGAGTACTTCCT<br>GACCTCGCCGGCGGGCCTGGTGAACATTGCCGAGAACTATGTGGGGCT<br>GCCGTTCGAGGAAGTTGCCTGA |
| *pobA* [c] | ATGAACCACGTACCAGTGGCAATTATTGGCGCAGGACCAGCAGGACTAA<br>CCCTCGCCCACCTCCTCCACCTTCAAGGTGTGGAATCAATCGTCTTTGAA<br>TCCCGCACCCGCAAGGACGTCGAAGAAACCGTCCGAGCAGGCATCCTG<br>GAACAAGGCACCCTGAATCTGATGCGCGAAACCGGAGTCGGCGCACGC<br>ATGGAAGCAGAAGCCGATCACGATGAAGCAATCGACATCTCCATCAACAA<br>TGAGCGCACCCGCATTCCGCTGACCGAACTCACCGGCCACAAGGTTGCG<br>ATCTACCCGCAGCACGAATACCTCAAAGATTTCATTGCCAAGCGCATCGA<br>AGATGGCGGCGAACTCCTTTTCACCACCACTGTTGATTCCGTAGAAAACT<br>ACGAAGGCGACCTCGCCAAGGTGACCTACACCGAAGCCGATGGTTCCTC<br>CACCACCATCACCGCCGACTACGTCATCGCAGCTGACGGCTCCAACTCC<br>CCTTACCGCAAGCTGATCACCGAAGACGGTGGCGTGCGCGCCCGCCAT |

| | |
|---|---|
| *pobA* [c] | GAATACCCTTACGCATGGTTCGGCATTTTGGTGGAAGCACCAAAAACCCA<br>AAAGGAACTCATCTACGCAACCCACCCTGAGGGCTTTGCGCTGATCTCC<br>ACCCGTACCGATGAAATCCAGCGCTACTACCTGCAGTGCAACCCTGACG<br>ACACCCCAGACATGTGGCCCGATGACCGCATTTGGGAACAGCTGCACCT<br>GCGTGCGGACTCCCCTGGCATCACCGTGTCTGAAGGGCGCATCTTTGAC<br>AAGGCCGTGCTGCGTTTCCGCTCCGCGGTCACCGAACCAATGCAAAAGG<br>GACGCCTCTTCCTTGCTGGCGATGCTGCACACACCGTGCCGCCAACCGG<br>AGCTAAGGGCCTCAACTTGGCTGTTGCCGATGTCTCAGTACTCGCGCCA<br>GCACTGGTTCGTGCCCTGAAGAAGAAGGACACCGGCTTGCTCGATAGCT<br>ACACCTCCCTGGCAGTCCCCCGCATCTGGAAAGCACAGCACTTCTCCTA<br>CTGGATGAGCTCCATGCTCCACGCAGTACCCGGCGAAGATCACTTTGCC<br>ACCCAGCGCCGATTCGCTGAATTGCGCTCCGTCCTAGAATCCCAATCCG<br>GCCAACGCTACCTCGCAGAGCAGTACGTTGGGCGCGACCTACCACGCTT<br>CGAGGTATAA |

[a] Synthesized gene was carried by pUC57 plasmid with Amp$^R$.

[b] The *pobA* gene was from *Pseudomonas putida*.

[c] The *pobA* gene was from *Corynebacterium glutamicum*.

**Table S10.** gRNA sequences used in this study.

| gRNA | Target | PAM | Strand [a] | spacer |
|---|---|---|---|---|
| gRNA-pcaI | *pcaI* | AGA | C | gctgcagacgcggcaggtga |
| gRNA-pcaJ | *pcaJ* | CGG | C | caatcagaaaacgggatgct |

[a] C stands for coding strand.

**Table S11.** The PobA used for phylogenetic analysis in this study.

| Organism | Accession number |
|---|---|
| *Roseovarius nubinhibens* | WP_009814201.1 |
| *Ruegeria pomeroyi* | WP_011241830.1 |
| *Dinoroseobacter shibae* | WP_012180136.1 |
| *Roseobacter denitrificans* | WP_011568529.1 |
| *Roseobacter litoralis* | WP_044025630.1 |
| *Corynebacterium glutamicum* | WP_011014104.1 |
| *Rhodococcus jostii* | WP_011595255.1 |
| *Enterobacter cloacae* | SAJ20715.1 |
| *Pseudomonas putida* | WP_010954394.1 |
| *Comamonas testosteroni* | WP_003055490.1 |

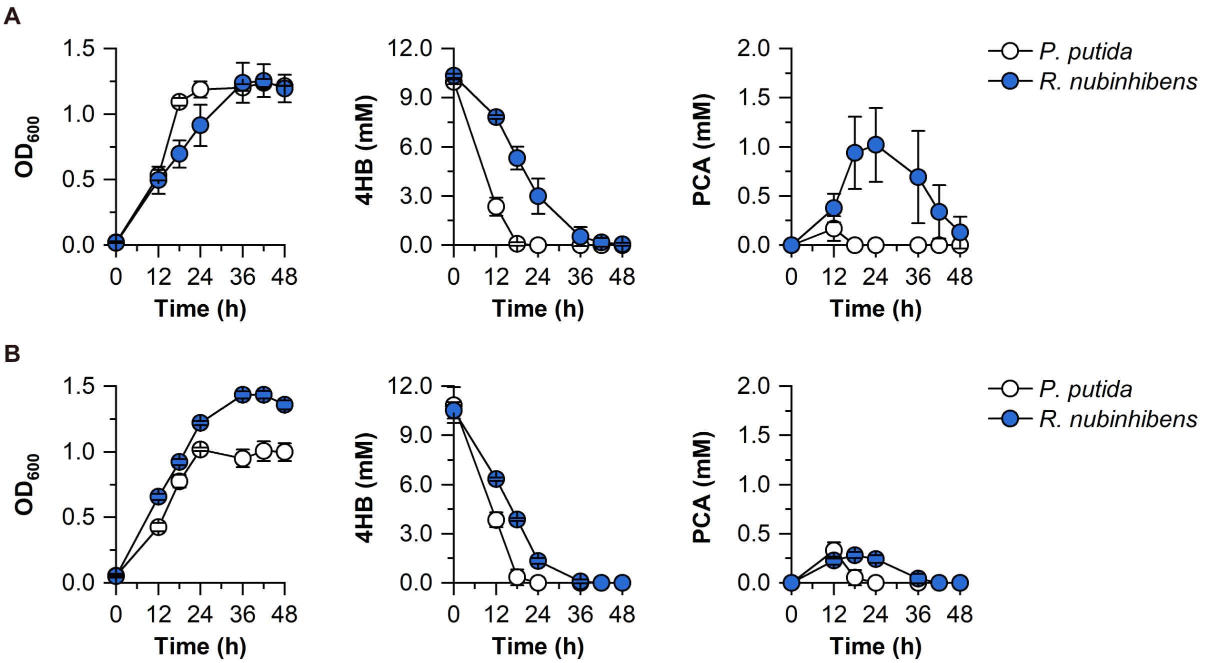


**Fig. S1. Performance of *R. nubinhibens* and *P. putida* in SBM. (A)** The cultivation experiments were conducted at pH 8.1 and **(B)** at pH 8.5. Experiments were carried out in triplicate and the error bars represented the standard deviations of the means of three biological replicates.

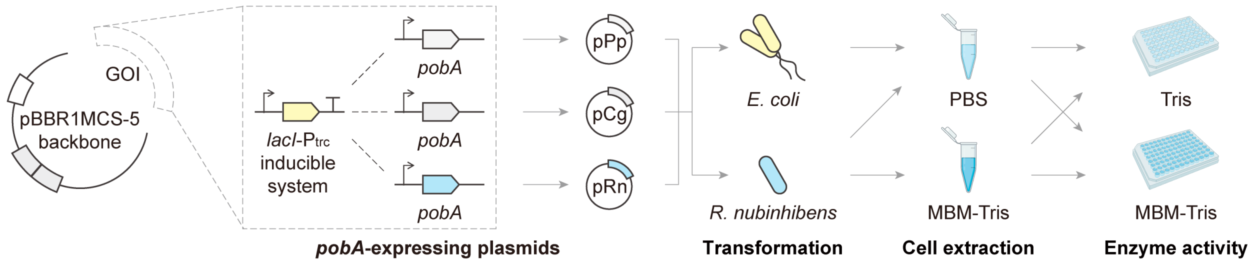


**Fig. S2. Schematic illustration of plasmid design and enzyme activity analysis.** Three *pobA* genes from *P. putida*, *C. glutamicum* and *R. nubinhibens* were cloned to pBBR1MCS-5 backbone and controlled by the *lacI*-$P_{trc}$ inducible system. The resulting plasmids were transformed to *E. coli* BW25113 and *R. nubinhibens*, respectively. For *E. coli*, crude cell extract was prepared in PBS and for *R. nubinhibens*, both in PBS and MBM-Tris. The enzyme activity was analyzed in Tris or MBM-Tris at different pHs. GOI, gene of interest; PBS, phosphate buffered saline; MBM, marine basal media; Tris, tris (hydroxymethyl) aminomethane. Created with BioRender.com.

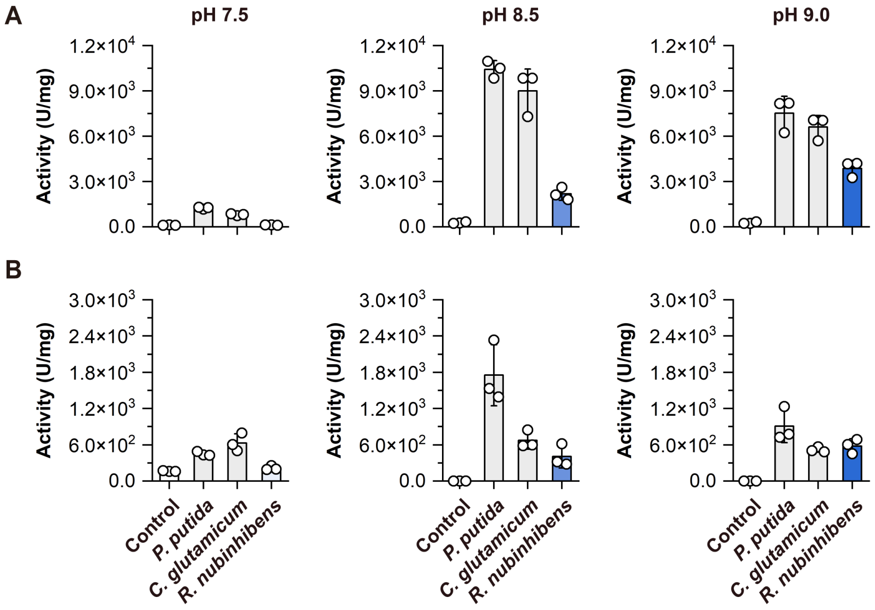


**Fig. S3. Enzyme activity of PobA. (A)** Enzyme activity of three PobA enzymes analyzed in Tris buffer and **(B)** MBM-Tris buffer. The *pobA* genes were expressed in *E. coli* and the enzymes were extracted in PBS. The experiments were conducted in triplicate, and the circles and error bars represent the individual values and standard deviations of three biological replicates, respectively.

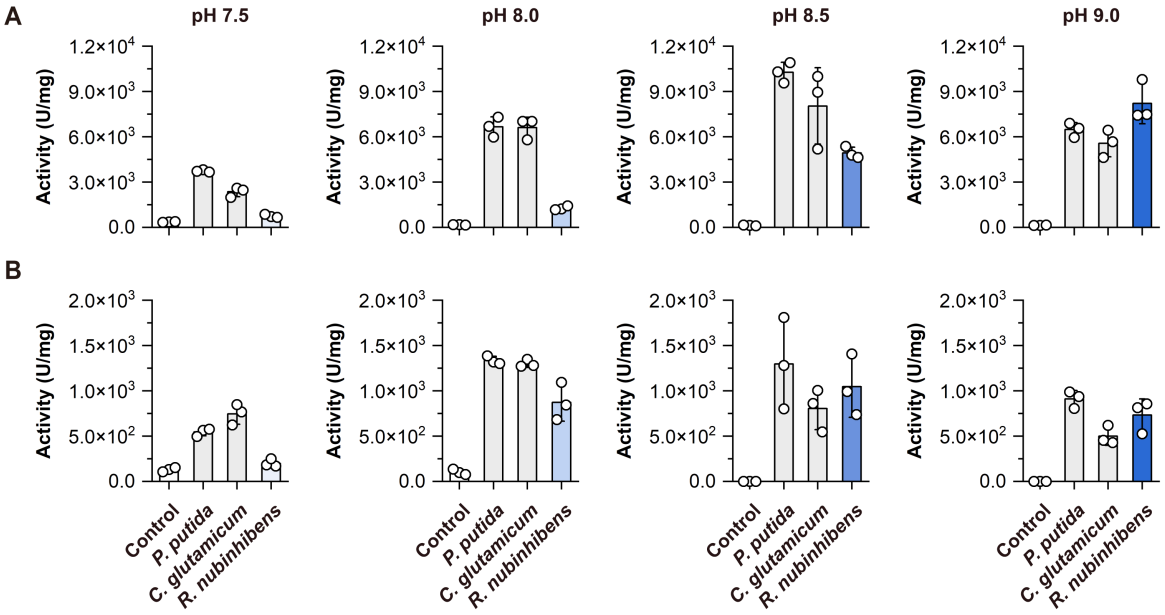


**Fig. S4. Enzyme activity of PobA. (A)** Enzyme activity of three PobA enzymes analyzed in Tris buffer and **(B)** MBM-Tris buffer. The *pobA* genes were expressed in *R. nubinhibens* and the enzymes were extracted in PBS. The experiments were conducted in triplicate, and the circles and error bars represent the individual values and standard deviations of three biological replicates, respectively.

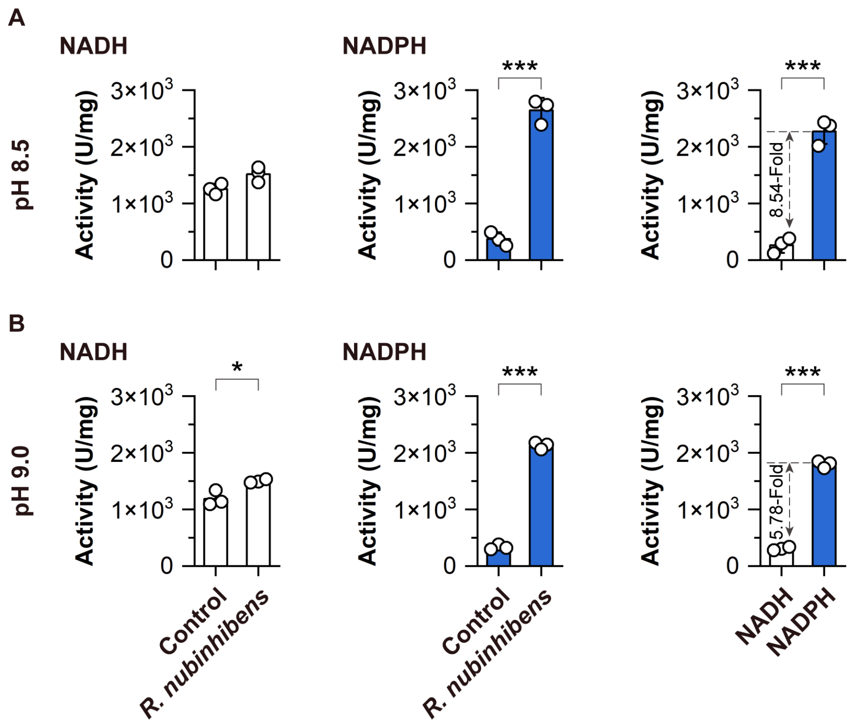


**Fig. S5. NAD(P)H specificity of PobA from *R. nubinhibens*. (A)** The enzyme activity analysis was conducted at pH 8.5 and **(B)** at pH 9.0. The experiments were conducted in triplicate, and the circles and error bars represent the individual values and standard deviations of three biological replicates, respectively. The differences were statistically evaluated by *t*-test (*, P<0.05; ***, P<0.001).

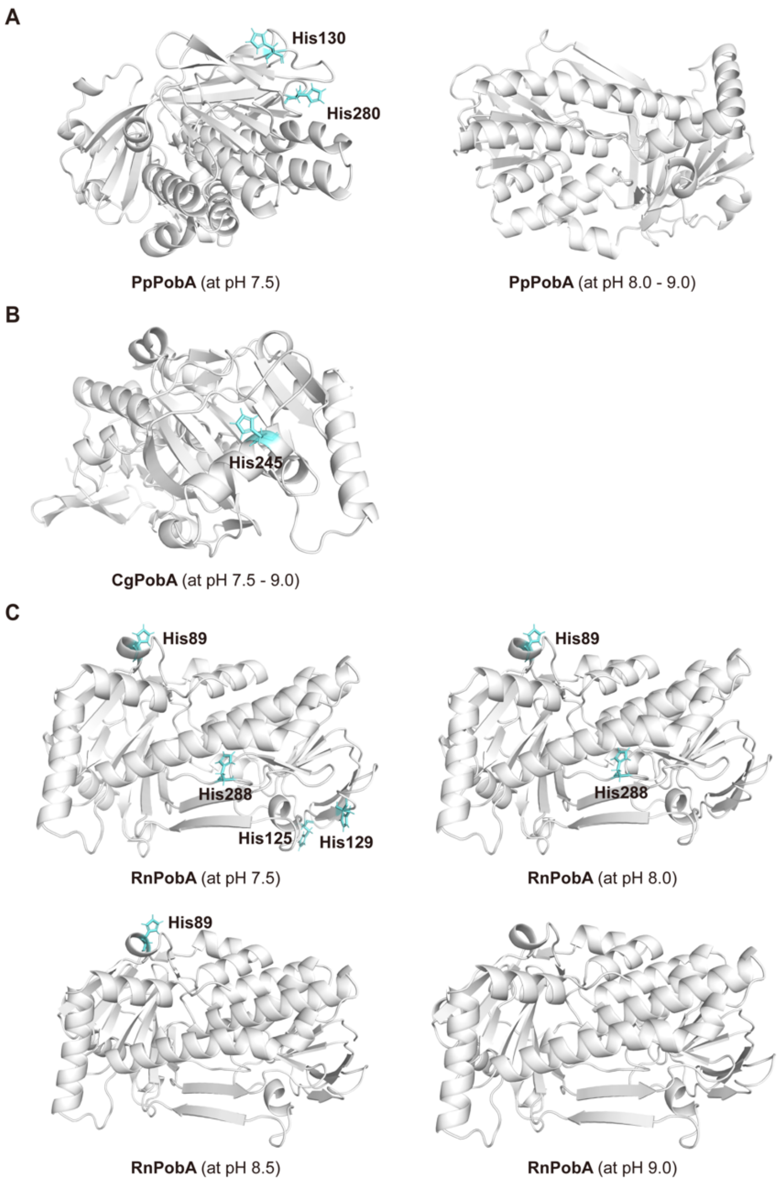


**Fig. S6. Structural heterogeneity of PobA. (A)** Structure of PpPobA, **(B)** CgPobA and **(C)** RnPobA at different pHs. The protonations of histidine are indicated in blue.

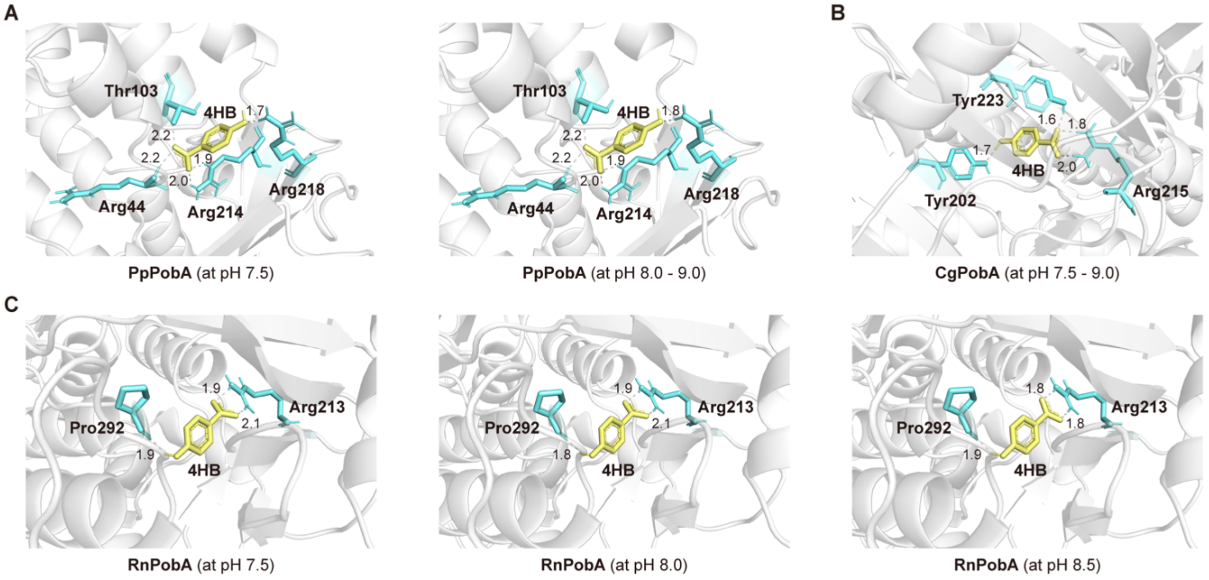


**Fig. S7. Molecular docking results of PobA with 4HB. (A)** Molecular docking results of PpPobA, **(B)** CgPobA and **(C)** RnPobA with 4HB at different pHs. The residues of PobA identified as the binding site to 4HB are indicated in blue and the hydrogen bonds are indicated in grey.

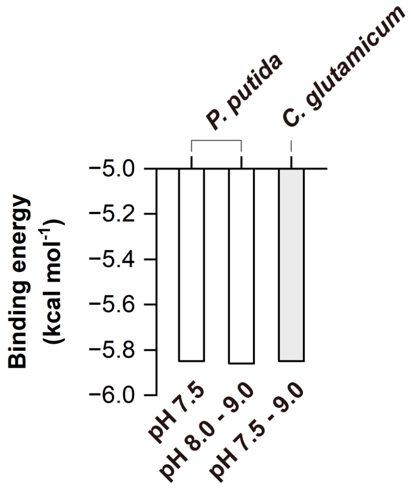


**Fig. S8. Binding energy of PobA with 4HB.** The binding energy of PpPobA with 4HB is indicated in white columns and that of CgPobA with 4HB is indicated in grey column.

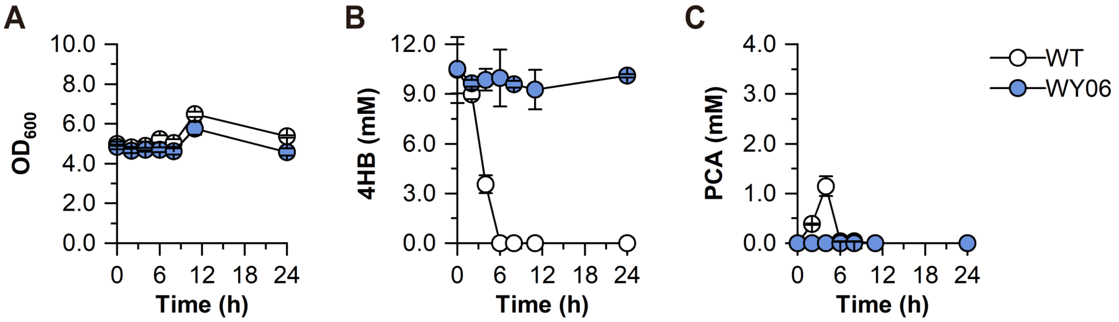


**Fig. S9. Biosynthetic performance of concentrated WY06 with the initial $OD_{600}$ at 5.0. (A)** Growth profile. **(B)** Consumption of 4HB. **(C)** Production of PCA. The experiments were conducted in triplicate and the error bars represent the standard deviations of three biological replicates, respectively.

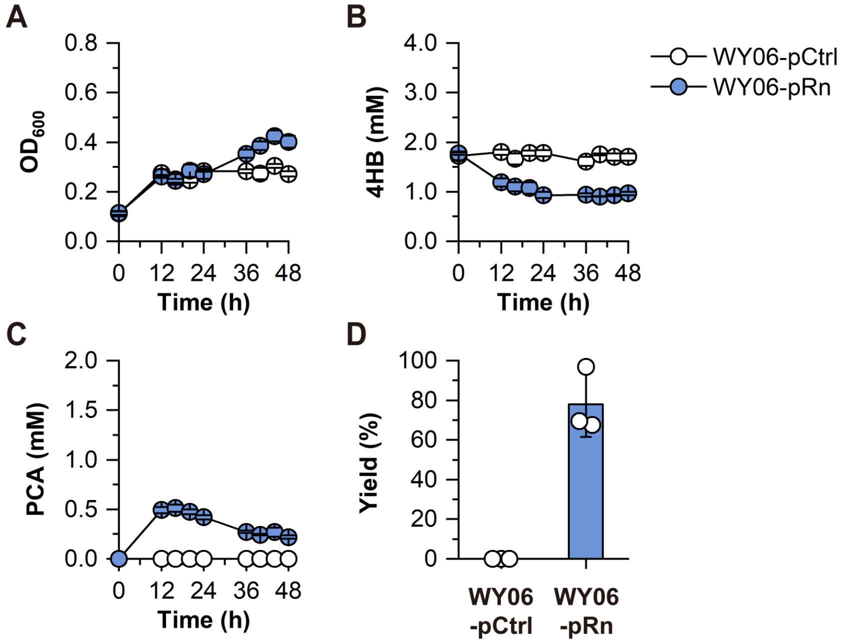


**Fig. S10. Biosynthetic performance of WY06-pRn with 2 mM 4HB. (A)** Growth profile. **(B)** Consumption of 4HB. **(C)** Production of PCA. **(D)** Molar yield. The experiments were conducted in triplicate, and the circles and error bars represent the individual values and standard deviations of three biological replicates, respectively.

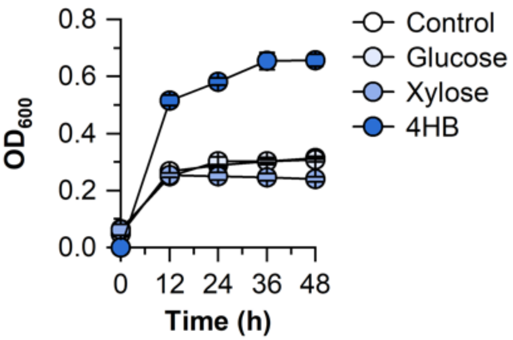


**Fig. S11. Growth profile of *R. nubinhibens*.** The strains were cultured in MBM with 5 g/L glucose, 5 g/L xylose or 10 mM 4HB. The Control group was cultivated in MBM without additional carbon source. Experiments were carried out in triplicate and the error bars represented the standard deviations of the means of three biological replicates.

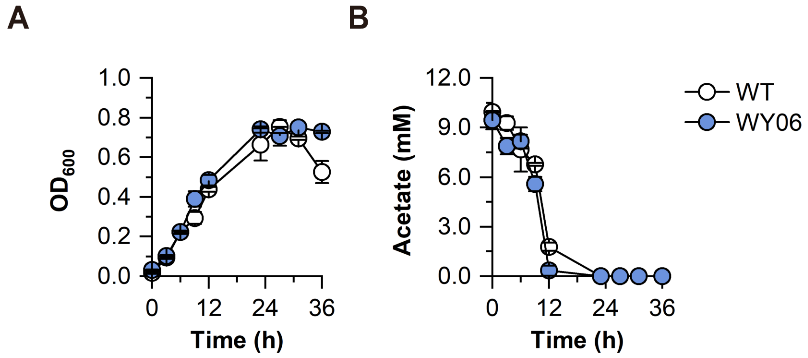


**Fig. S12. Growth of *R. nubinhibens* on acetate. (A)** Growth profile. **(B)** Consumption of acetate. The experiments were conducted in triplicate and the error bars represent the standard deviations of three biological replicates, respectively.

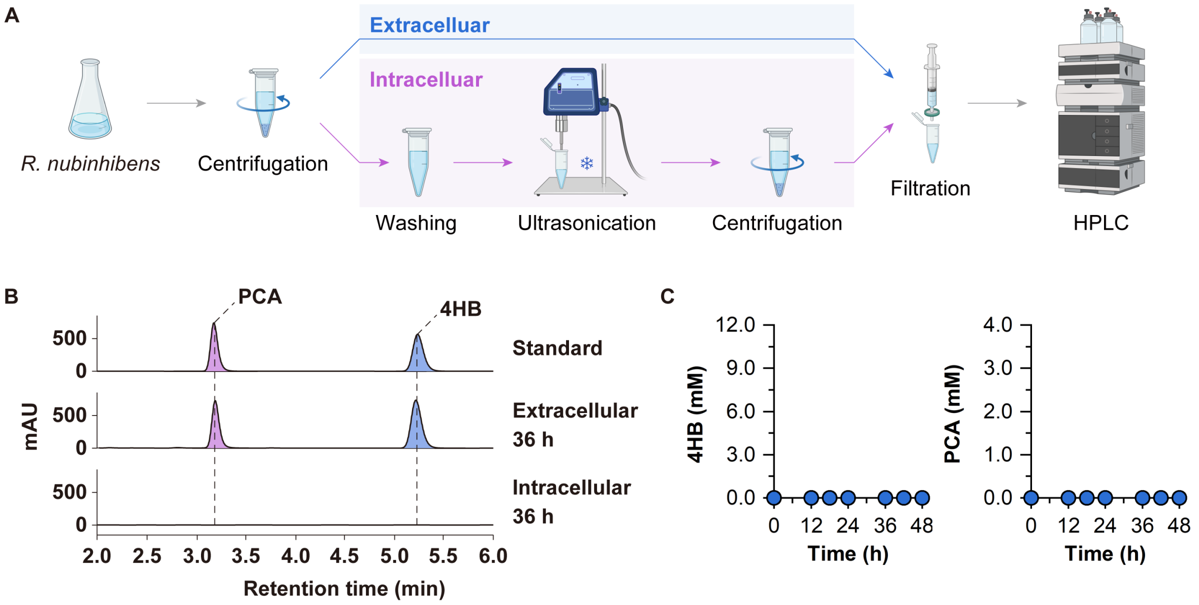


**Fig. S13. Evaluation of the intracellular PCA. (A)** Schematic illustration of the intracellular and extracellular PCA analysis. **(B)** HPLC profiles of the standard and the extracellular and intracellular samples at 36 h. **(C)** Results of the intracellular 4HB and PCA. Panel **A** Created with BioRender.com.

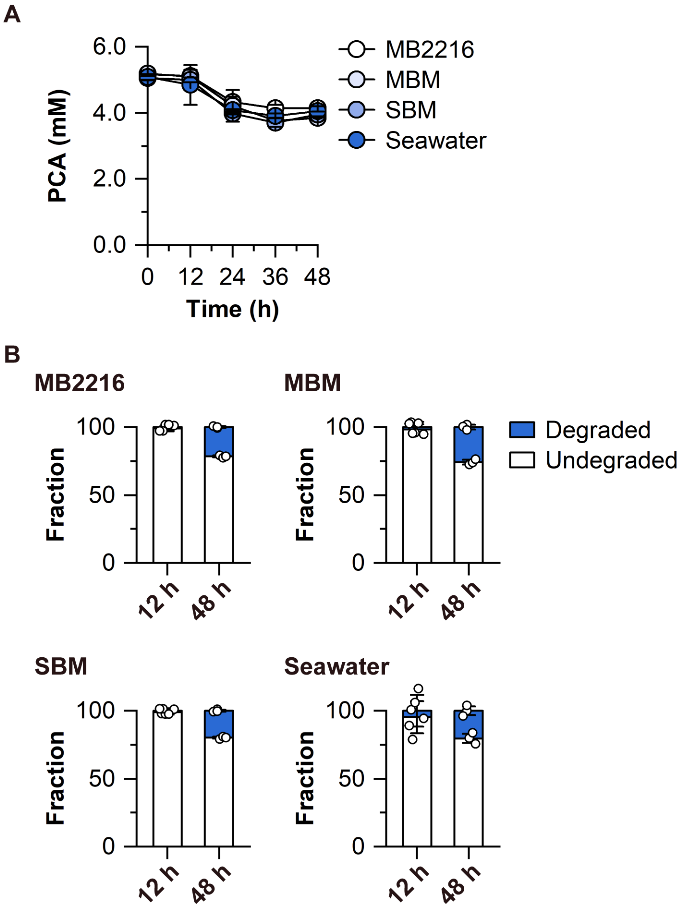


**Fig. S14. Stability of PCA in different media. (A)** PCA concentration in 48 h. **(B)** Fraction of degraded and undegraded PCA at 12 h and 48 h. MB2216, the rich medium. MBM, the synthetic minimal medium. SBM, the seawater-based minimal medium. Seawater, after filtration. Experiments were carried out in triplicate and the error bars represented the standard deviations of the means of three biological replicates.

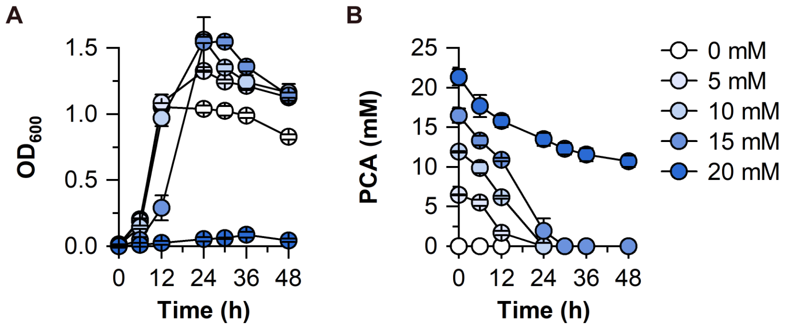


**Fig. S15. Tolerance of *R. nubinhibens* to PCA. (A)** Growth profile. **(B)** PCA concentration. Experiments were carried out in triplicate and the error bars represented the standard deviations of the means of three biological replicates.

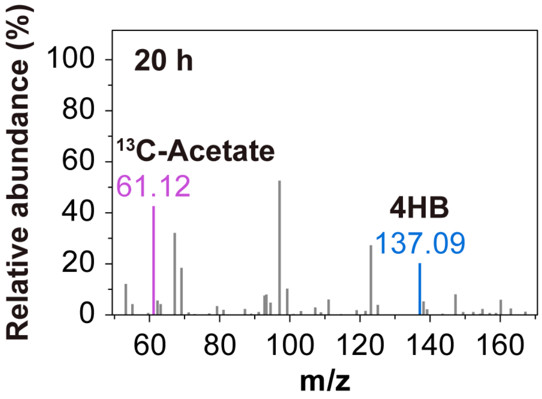


**Fig. S16. MS analysis of samples of the control strain WY10-pCtrl.** The $^{13}$C-labeled acetate is indicated in purple and the non-labeled 4HB is indicated in blue.

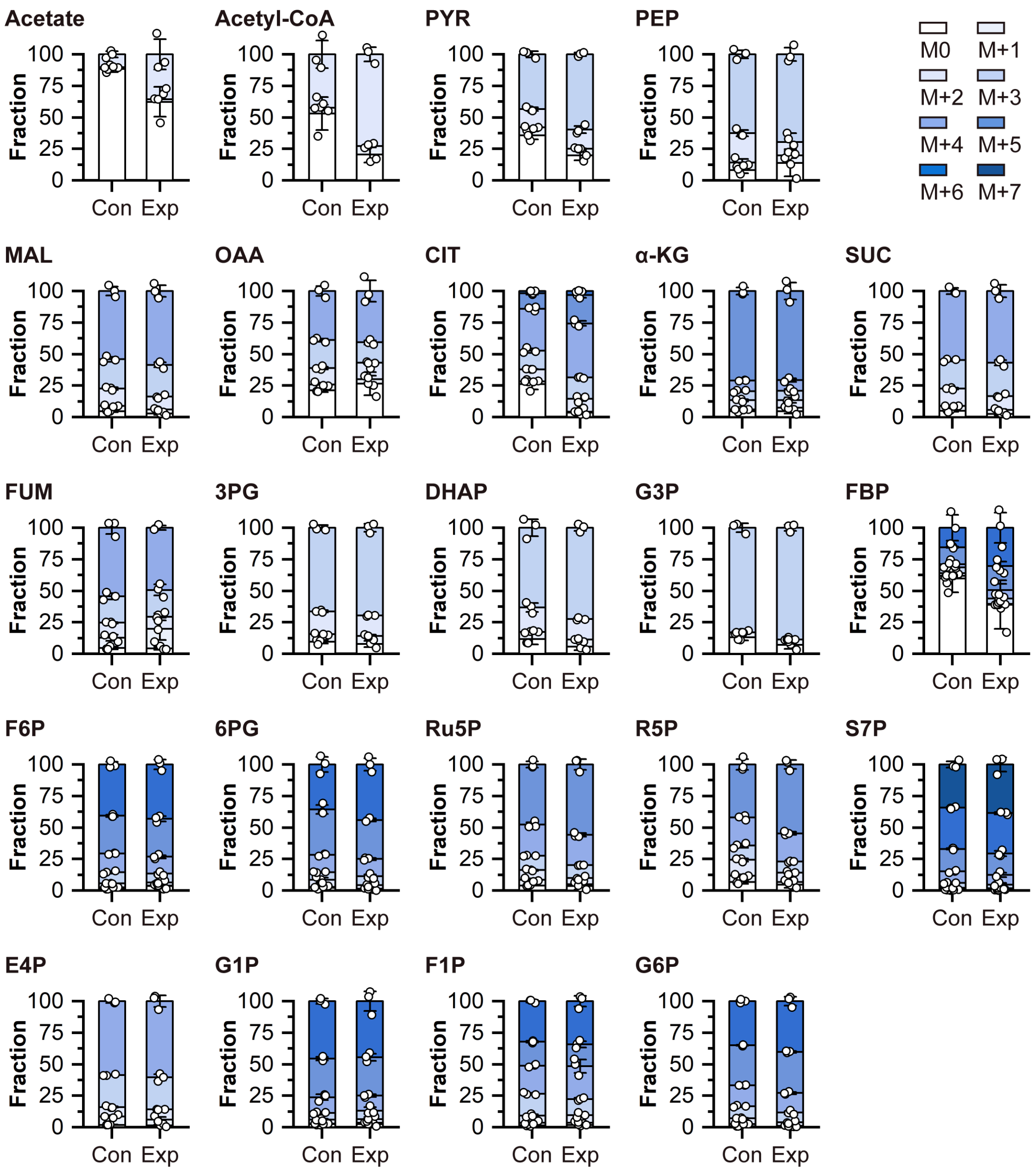


**Fig. S17. Isotopic labeling patterns of key metabolites in WY10-pCtrl (Con) and WY10-pRn (Exp).** Both strains were cultivated with 4HB and $^{13}$C-acetate. M0 represents the fragments without labeled carbon and M + *n* represents the fragments with *n* $^{13}$C-labeled carbon. **Abbreviations:** PYR, pyruvate; PEP, phosphoenolpyruvate; MAL, malate; OAA, oxaloacetate; CIT, citrate; α-KG, α-ketoglutarate; SUC, succinate; FUM, fumarate; 3PG, 3-phosphoglycerate; DHAP, dihydroxyacetone phosphate; G3P, glyceraldehyde-3-phosphate; FBP, fructose-1,6-bisphosphate; F6P, fructose-6-phosphate; 6PG, 6-phosphogluconate; Ru5P, ribulose-5-

phosphate; R5P, ribose-5-phosphate; S7P, sedoheptulose-7-phosphate; E4P, erythrose-4-phosphate; G1P, glucose-1-phosphate; F1P, fructose-1-phosphate; G6P, glucose-6-phosphate.

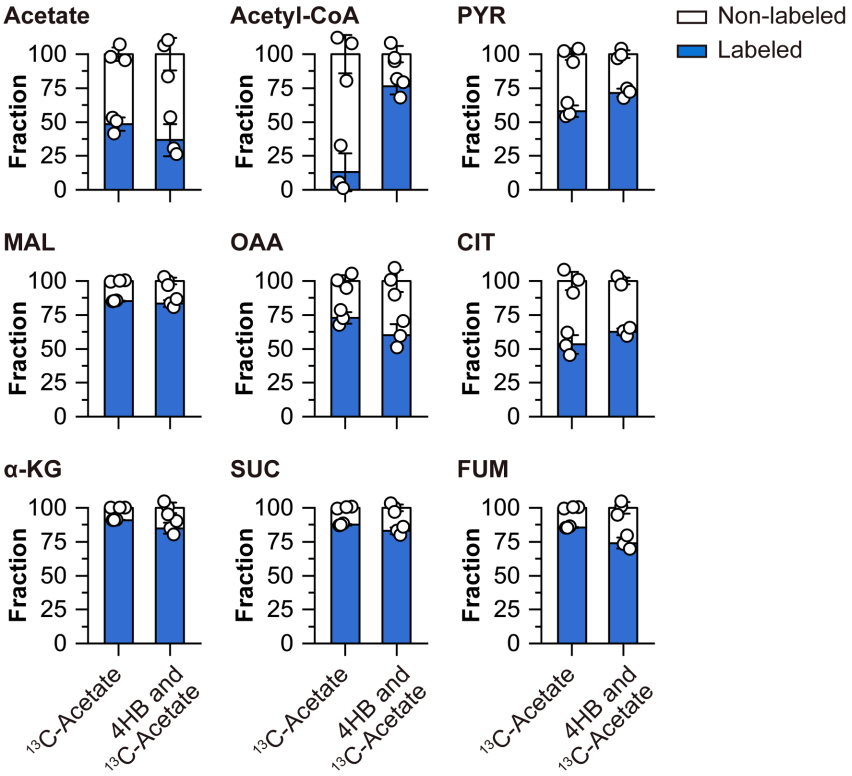


**Fig. S18. Isotopic labeling patterns of key metabolites in WY10-pRn.** The strain was cultivated with only $^{13}$C-acetate, or with both 4HB and $^{13}$C-acetate. **Abbreviations:** PYR, pyruvate; MAL, malate; OAA, oxaloacetate; CIT, citrate; α-KG, α-ketoglutarate; SUC, succinate; FUM, fumarate.

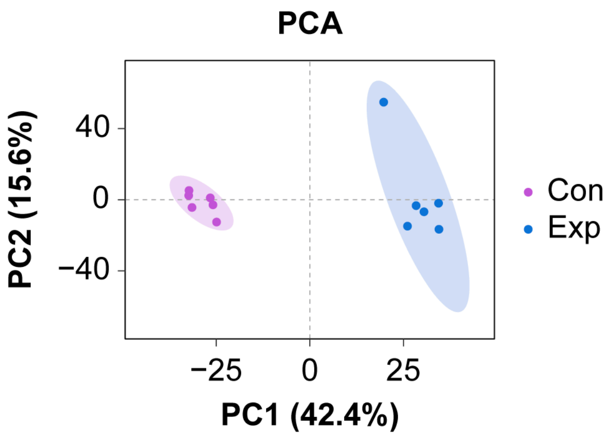


**Fig. S19. Principal component analysis (PCA).** Con and Exp represent the control strain WY10-pCtrl and the reprogrammed strain WY10-pRn, respectively.

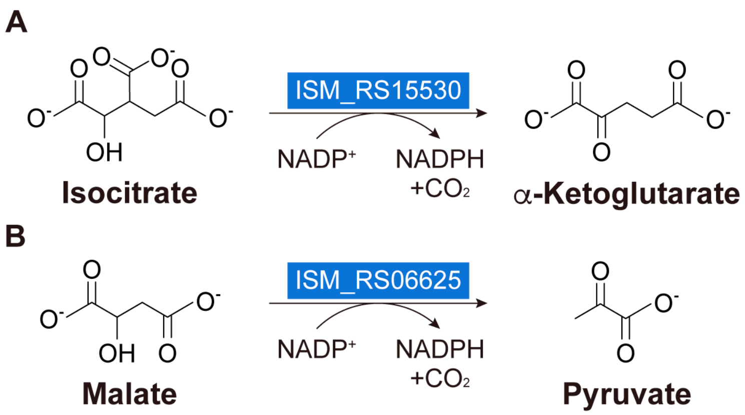


**Fig. S20. Reaction with NADPH generation. (A)** Conversion from isocitrate to α-ketoglutarate. **(B)** Conversion from malate to pyruvate.

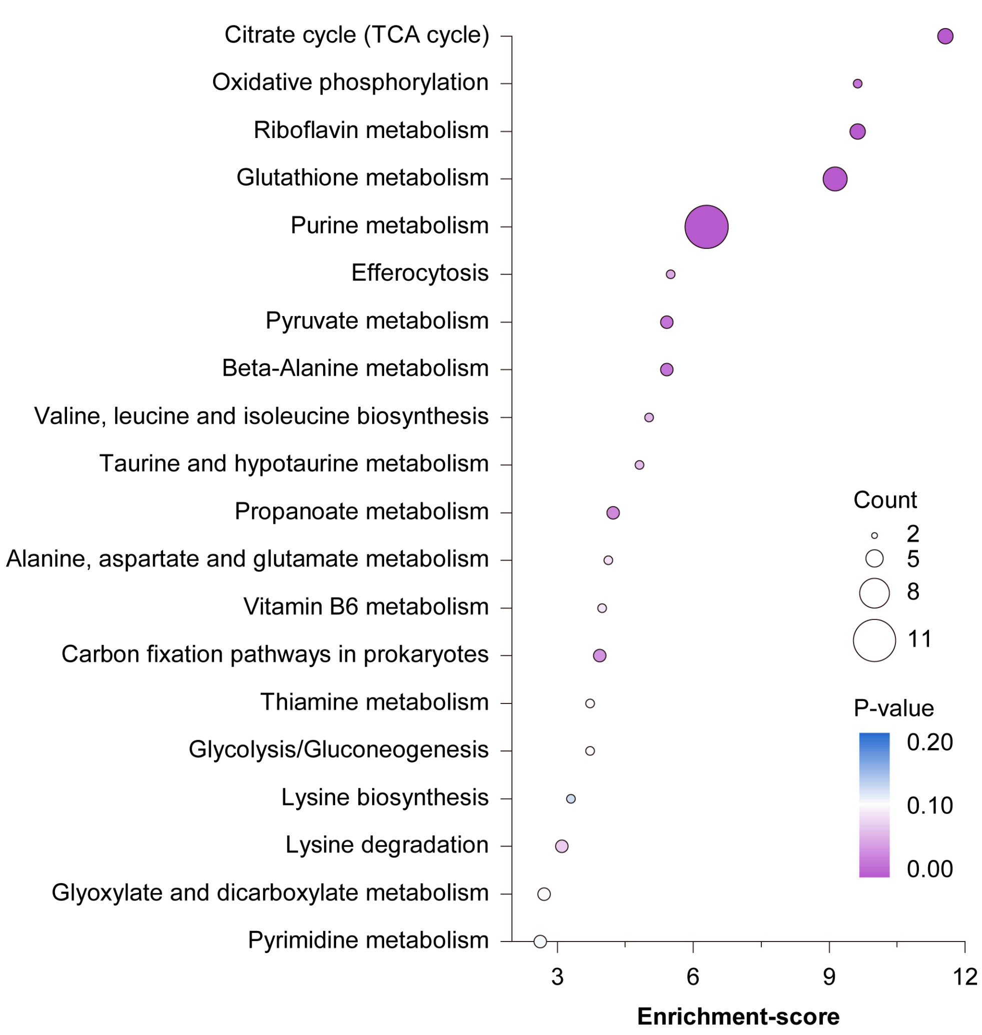


**Fig. S21. Significantly enriched KEGG pathways.** The size of circles represents the count of differential metabolites and the color represents the P-value.

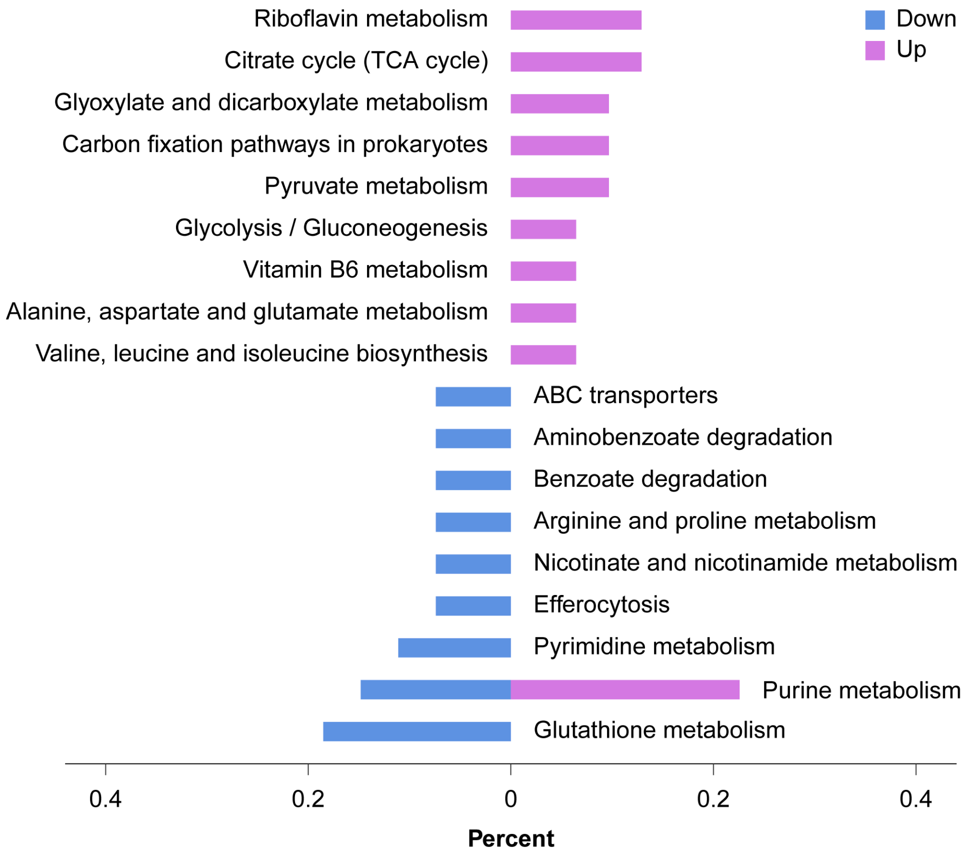


**Fig. S22. Comparison of upregulation and downregulation in enriched pathways.** The purple and blue bars represent upregulation and downregulation, respectively.

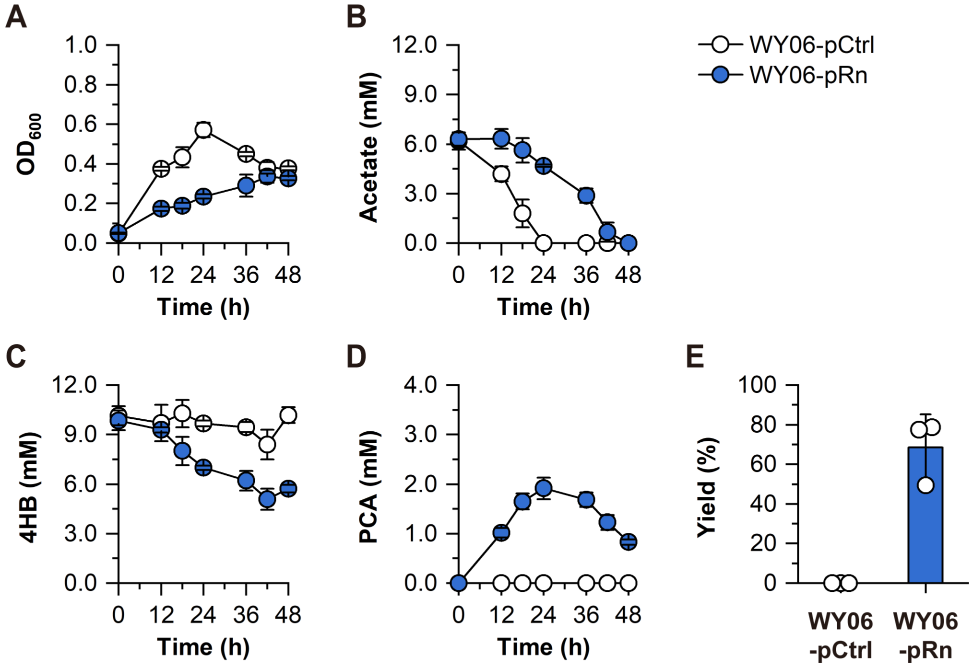


**Fig. S23. Biosynthetic performance of WY06-pRn with seawater as the water source. (A)** Growth profile. **(B)** Consumption of acetate. **(C)** Consumption of 4HB. **(D)** Production of PCA. **(E)** Molar yield. The experiments were conducted in triplicate, and the circles and error bars represent the individual values and standard deviations of three biological replicates, respectively.

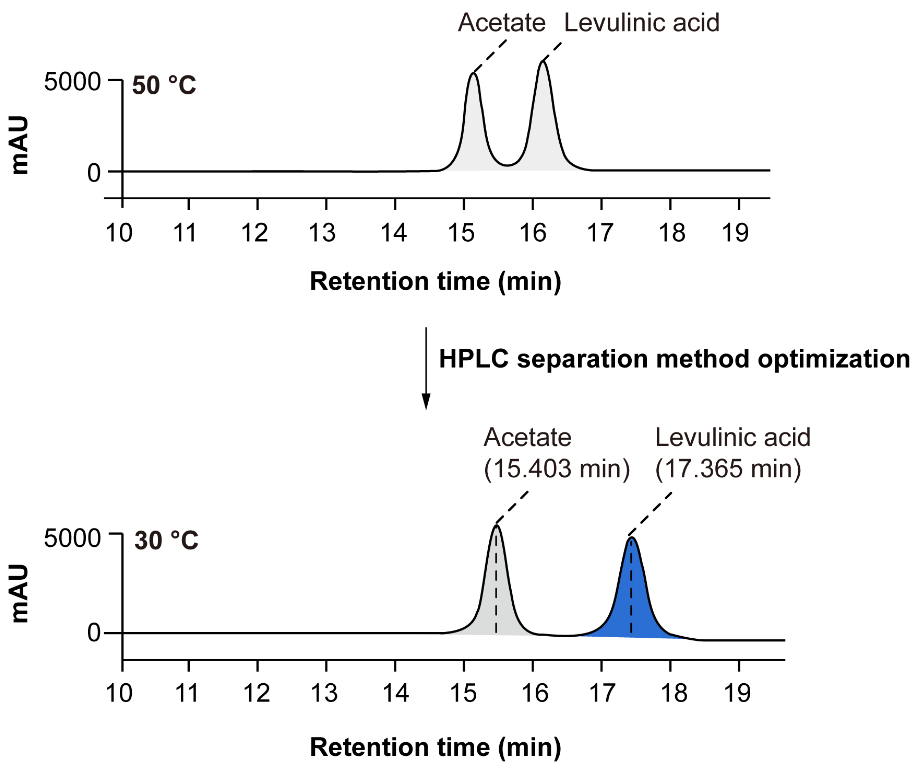


**Fig. S24. Optimization of HPLC separation of a mixed acetate and levulinic acid standard by decreasing temperature of column and detector.**
